# Ad26.COV2.S-elicited immunity protects against G614 spike variant SARS-CoV-2 infection in Syrian hamsters and does not enhance respiratory disease in challenged animals with breakthrough infection after sub-optimal vaccine dosing

**DOI:** 10.1101/2021.01.08.425915

**Authors:** Joan E.M. van der Lubbe, Sietske K. Rosendahl Huber, Aneesh Vijayan, Liesbeth Dekking, Ella van Huizen, Jessica Vreugdenhil, Ying Choi, Miranda R.M. Baert, Karin Feddes-de Boer, Ana Izquierdo Gil, Marjolein van Heerden, Tim J. Dalebout, Sebenzile K. Myeni, Marjolein Kikkert, Eric J. Snijder, Leon de Waal, Koert J. Stittelaar, Jeroen T.B.M. Tolboom, Jan Serroyen, Leacky Muchene, Leslie van der Fits, Lucy Rutten, Johannes P.M. Langedijk, Dan H. Barouch, Hanneke Schuitemaker, Roland C. Zahn, Frank Wegmann

## Abstract

Previously we have shown that a single dose of recombinant adenovirus serotype 26 (Ad26) vaccine expressing a prefusion stabilized SARS-CoV-2 spike antigen (Ad26.COV2.S) is immunogenic and provides protection in Syrian hamster and non-human primate SARS-CoV-2 infection models. Here, we investigated the immunogenicity, protective efficacy and potential for vaccine-associated enhanced respiratory disease (VAERD) mediated by Ad26.COV2.S in a moderate disease Syrian hamster challenge model, using the currently most prevalent G614 spike SARS-CoV-2 variant. Vaccine doses of 1×10^9^ vp and 1×10^10^ vp elicited substantial neutralizing antibodies titers and completely protected over 80% of SARS-CoV-2 inoculated Syrian hamsters from lung infection and pneumonia but not upper respiratory tract infection. A second vaccine dose further increased neutralizing antibody titers which was associated with decreased infectious viral load in the upper respiratory tract after SARS-CoV-2 challenge. Suboptimal non-protective immune responses elicited by low-dose A26.COV2.S vaccination did not exacerbate respiratory disease in SARS-CoV-2-inoculated Syrian hamsters with breakthrough infection. In addition, dosing down the vaccine allowed to establish that binding and neutralizing antibody titers correlate with lower respiratory tract protection probability. Overall, these pre-clinical data confirm efficacy of a 1-dose vaccine regimen with Ad26.COV2.S in this G614 spike SARS-CoV-2 virus variant Syrian hamster model, show the added benefit of a second vaccine dose, and demonstrate that there are no signs of VAERD under conditions of suboptimal immunity.

## INTRODUCTION

Severe Acute Respiratory Syndrome Coronavirus 2 (SARS-CoV-2), the etiological agent of coronavirus disease 2019 (COVID-19), is responsible for an unprecedented crisis in the world^1^. While physical measures such as social distancing and deployment of face masks are being employed to reduce the spread of the virus, safe and effective vaccines are crucial to contain this pandemic.

We recently demonstrated that a single dose of adenovirus serotype 26 (Ad26) vaccine expressing a pre-fusion stabilized spike antigen (Ad26.CoV2.S) is immunogenic in animals and humans^2–5^ and protected rhesus macaques against challenge with SARS-CoV-2^2^. Protection correlated with SARS-CoV-2 binding and neutralizing antibodies, in agreement with findings with other COVID-19 vaccine candidates^6–8^.

We have also demonstrated vaccine mediated protection against SARS-CoV-2 challenge in Syrian hamsters. In this animal model, SARS-CoV-2 infection is characterized by severe clinical disease including body weight loss and respiratory tract histopathology upon high-dose intranasal challenge^3,9–11^, mimicking findings in humans where viral inoculum size has been correlated with disease severity^12–15^. In those studies, challenge in Syrian hamsters was performed with SARS-CoV-2 USA-WA1/2020 which has a 100% homologous spike sequence to the Ad26.COV2.S vaccine antigen^3,4^. Since then, SARS-CoV-2 variants with a D614G spike substitution have become most prevalent^16,17^, albeit recently new strains with additional mutations in spike further distant from the original Wuhan strain are emerging and became more widespread^18–20^. In Syrian hamsters, comparison of a SARS-CoV-2 strain with and without the D614G substitution in the spike protein indicated that the G614 variant produced higher infectious viral titers in the upper respiratory tract and increases competitive fitness ^21,22^.

A potential concern of coronavirus vaccines is that they may predispose for disease enhancement after breakthrough infection, by eliciting only low- or non-neutralizing antibodies in combination with a Th2 skewed cellular response^14,23–25^. Vaccine-associated enhanced respiratory disease (VAERD) has been described for vaccine candidates for SARS-CoV and MERS-CoV but only in some animal models^26–30^. No human studies for these coronavirus vaccines have been reported that would allow a confirmation of the predictive value of these animal models. To our knowledge, there is no evidence of VAERD in nonclinical studies with SARS-CoV-2 vaccines available to date. Furthermore, clinical studies with SARS-CoV-2 vaccines, including the large-scale Phase 3 studies that are currently ongoing, have so far not reported any VAERD events.

The Ad26 vaccine vector is being used in a licensed Ebola virus vaccine^31,32^ and in multiple candidate vaccine programs, and has uniformly induced potently neutralizing antibody- and cellular immune responses with a clear Th1 skewing in non-clinical and clinical studies^33–36^. Similarly, Ad26.COV2.S induced neutralizing antibodies and a Th1 skewed cellular immune response in mice^4^, NHP^2^ and humans^5^, however, additional animal studies at suboptimal immunity to allow breakthrough infection are considered important to address the potential risk of predisposition for VAERD by Ad26.COV2.S.

Here we established an additional SARS-CoV-2 hamster challenge model and utilized it to confirm immunogenicity of Ad26.COV2.S and to verify its protective efficacy against intranasal infection with a heterologous G614 spike variant of SARS-CoV-2 (BetaCoV/Munich/BavPat1/2020). Furthermore, we established immunogenicity and efficacy of 2-dose Ad26.COV2.S regimens, confirmed correlates of protection and assessed VAERD by monitoring clinical, virological and histopathological signs of disease enhancements in hamsters receiving a sub-optimal dose of Ad26.COV2.S that did not protect against viral replication upon SARS-CoV-2 challenge.

## RESULTS

### Establishment of a G614 spike variant SARS-CoV-2 Syrian hamster challenge model

To assess vaccine immunogenicity, efficacy, and VAERD in hamsters, we established an Syrian hamster challenge model based on a SARS-CoV-2 strain with the D614G substitution in the spike protein. Male animals (n=12 per inoculation dose level) were inoculated with SARS-CoV-2 BetaCoV/Munich/BavPat1/2020 (containing a D614G substitution in the S1 fragment) at dose levels 10^2^, 10^3.3^, 10^4.6^ and 10^5.9^ 50% tissue culture infective dose (TCID_50_) administered by the intranasal route. Daily throat swabs were taken, and necropsies were performed 2, 3, 4, and 7 days post inoculation (dpi) (n=3 per timepoint), to monitor viral load in throat swabs, in lung and nose tissue, and to study respiratory tract pathology. As shown in Fig 1A and B, lung and nose tissue viral load assessment revealed high titers of replication-competent virus as measured by TCID_50_ in all inoculated animals at two dpi, independent of the size of the inoculum. The observed lung and nose viral load kinetics after two dpi were comparable across all tested inoculum quantities. Hamsters inoculated with 10^3.3^, 10^4.6^ and 10^5.9^ TCID_50_ showed highest viral loads in throat swabs at one dpi (Fig 1C), after which viral loads decreased to below the limit of detection by 4 to 5 dpi. By contrast, inoculation with the lowest SARS-CoV-2 dose of 10^2^ TCID_50_ resulted in an increase in infectious viral load in throat swabs from 1 to 2 dpi, suggesting viral replication, after which the viral load decreased to below the limit of detection by day 4 post inoculation.

**Figure 1.**
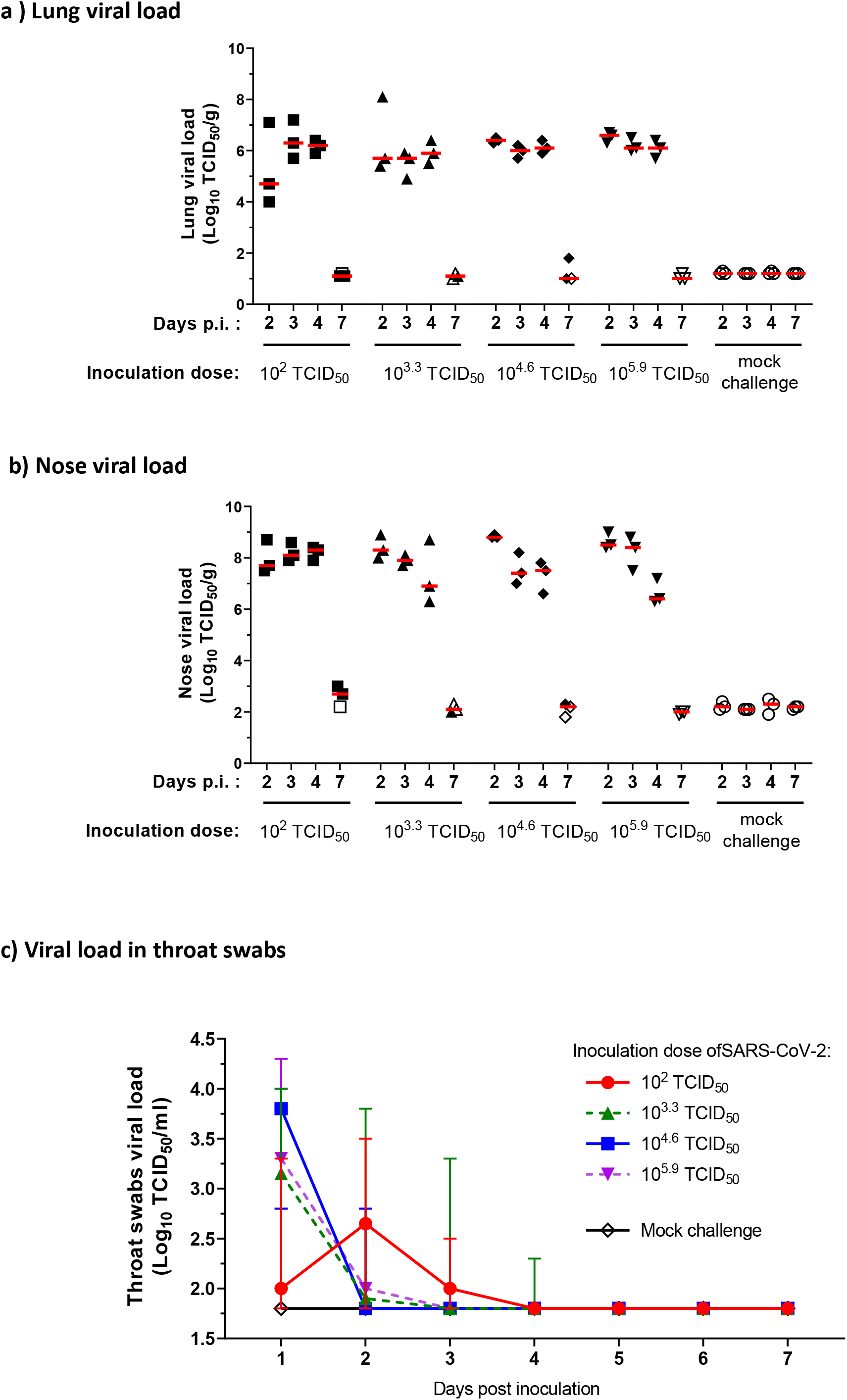
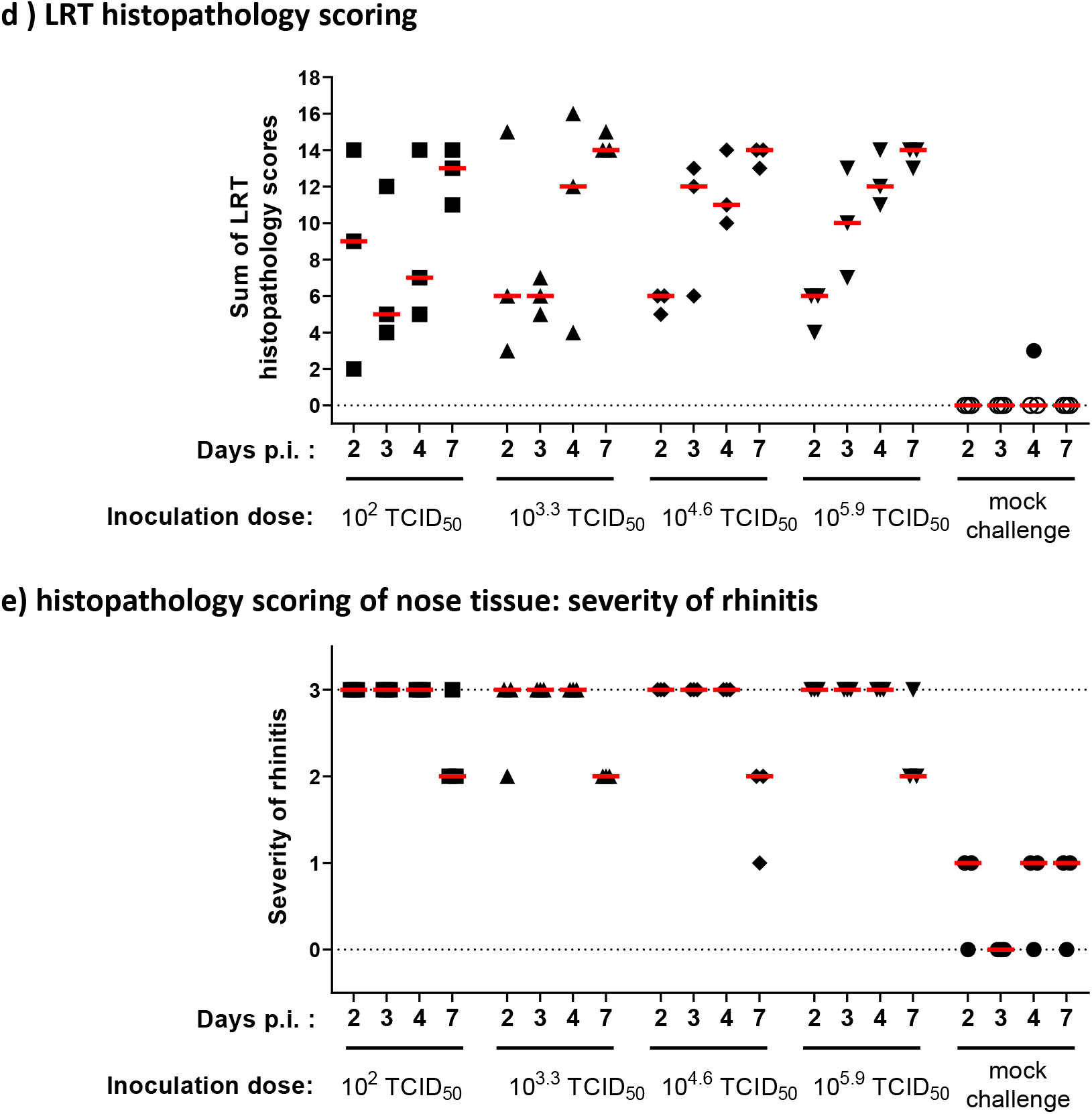
Titration of SARS-CoV-2 challenge dose and characterization of histopathology in Syrian hamsters. Syrian hamsters (N = 12 per group), were inoculated intra-nasally with 10^2^, 10^3.3^, 10^4.6^ or 10^5.9^ TCID_50_ SARS-CoV-2 BetaCoV/Munich/BavPat1/2020, or mock-inoculated with Vero E6 cell-supernatant. Daily throat swabs were taken, and 2, 3, 4, and 7 days p.i., 3 hamsters per group were sacrificed and nose and lung tissue collected for virological analysis and histopathology. Replication competent viral load in **a)** lung tissue, **b)** nose tissue, and **c)** throat swabs, was determined by TCID_50_ assay on Vero E6 cells. LLOD was calculated per animal per gram or milliliter of tissue, and animals with a response at or below the LLOD are shown as open symbols. **d)** Lung tissue was analyzed and scored for presence and severity of alveolitis, alveolar damage, alveolar edema, alveolar hemorrhage, type II pneumocyte hyperplasia, bronchitis, bronchiolitis, peribronchial and perivascular cuffing. Sum of scores are presented as sum of LRT disease parameters (potential range: 0 – 24). **e) N** ose tissue was analyzed and scored for severity of rhinitis on a scale from 0 to 3. Dotted lines indicate the minimal and maximal scores of histopathology. Median responses per group are indicated with horizontal lines, error bars in panel c indicate the range. p.i. = post inoculation; LLOD = lower limit of detection; LRT = lower respiratory tract; N = number of animals; TCID_50_/g = 50% tissue culture infective dose per gram tissue; TCID_50_/ml = 50% tissue culture infective dose per milliliter sample; vp = virus particles.

Histological analysis after challenge with 10^2^ TCID_50_ showed abundant presence of SARS-CoV-2 nucleocapsid protein (SARS-CoV-2 NP) by immunohistochemistry in areas of severe inflammation, characterized by multifocal moderate to severe degeneration and necrosis of upper and lower respiratory tract epithelial cells (Supp Fig 1). Compared with the higher challenge doses, 10^2^ TCID_50_ induced a comparable extent and severity of inflammation and damage throughout the respiratory tract, as determined by blinded semi-quantitative scoring (Fig 1D and E), with marginally lower lung histopathology scores at lower dose levels. Taken together, these results demonstrate that a low dose challenge inoculum induced a comparable viral load and disease pathology compared with higher viral dose challenges. For subsequent experiments we selected a 10^2^ TCID_50_ challenge dose associated with moderate disease based on histopathology findings, to allow assessment of the occurrence of more severe disease in this model and a theoretical risk for VAERD could be addressed. A 4-day follow up time after challenge was chosen as the most optimal time point to simultaneously evaluate lung tissue viral load and histopathology.

### Immunogenicity of Ad26.COV2.S in Syrian hamsters

Immunogenicity and protective efficacy of our Ad26.COV2.S vaccine candidate was assessed in the newly established challenge model described above. For comparison, two prototype Ad26 based vaccines were used expressing a membrane bound full length wild-type spike protein (Ad26.S) or a soluble pre-fusion stabilized spike protein with a C-terminal foldon replacing its transmembrane domain (Ad26.dTM.PP) ^2–4^. Male hamsters were immunized with either 10^9^ or 10^10^ viral particles (VPs) of Ad26.COV2.S and the two prototype vaccines. For each dose level, immunogenicity was assessed at various timepoints after a single immunization, and after a second homologous dose given 4 weeks later. Animals were challenged 4 weeks after one or two immunizations (Fig 2A). At week 4, Ad26.COV2.S elicited the highest neutralizing antibody titers and frequency of responding animals across dose levels (median titer 10^9^ VP 22.6, 10^10^ VP 38.6; 12/12 responders) compared with Ad26.S (median titer 10^9^ VP 8.5, 10^10^ VP 9.7; 8/12 responders, p<0.001) and Ad26.dTM.PP (median titer 10^9^ VP 8.5, 10^10^ VP 16.0; 8/12 responders, p<0.001) (Fig 2B). A second dose, irrespective of vaccine used, increased neutralization titers (week 8; Fig 2C). The Ad26.COV2.S vaccine was most immunogenic also after two doses, with a median neutralization titer of 128 after two doses of 10^9^ VP and 219 after two doses of 10^10^ VP, compared with neutralization titers of 32 and 55 for Ad26.S (p=0.003), and 16 and 61 for A26.dTM.PP (p=0.002), respectively.

**Figure 2.**
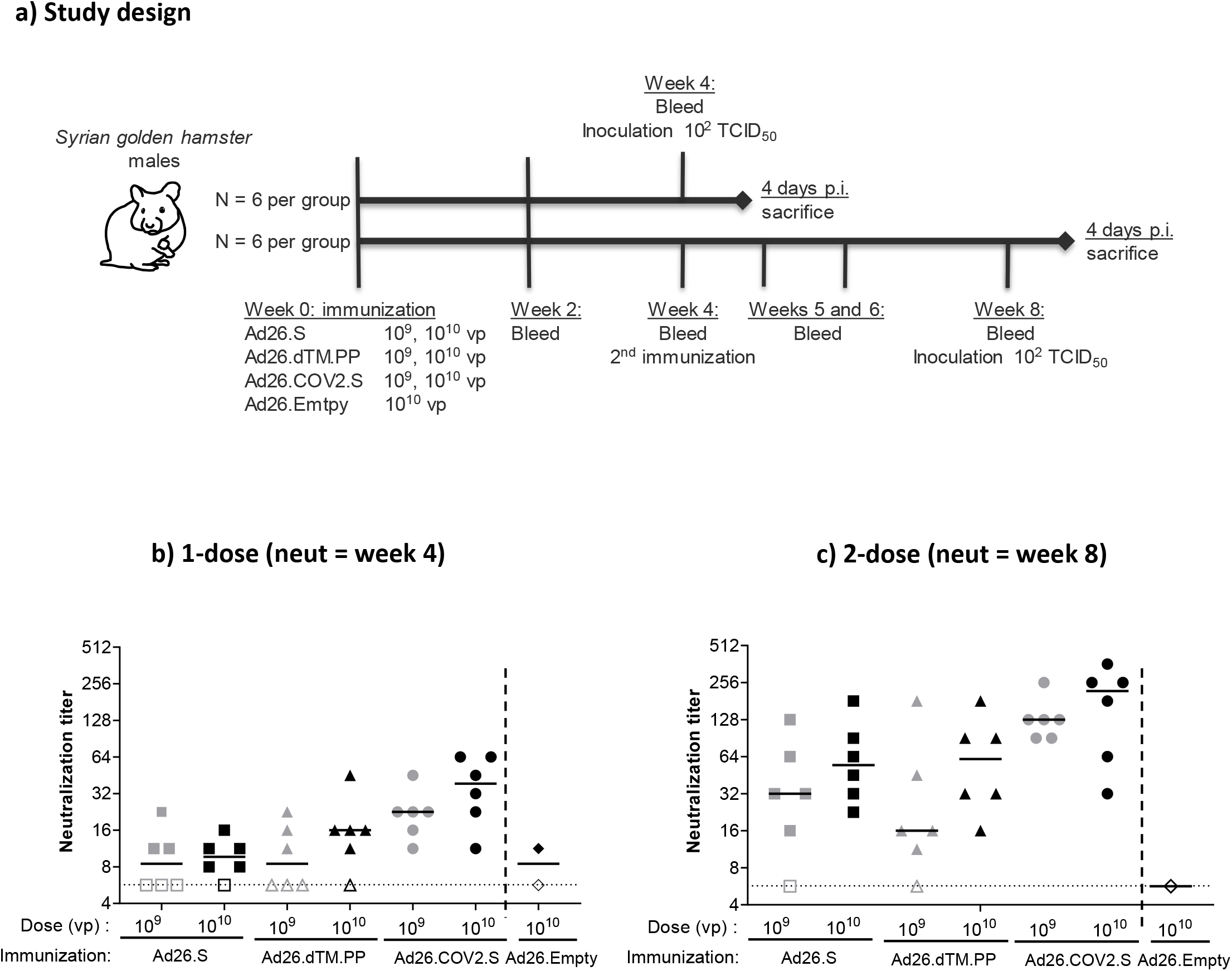
SARS-CoV-2 neutralizing antibody response elicited by 1- and 2-dose Ad26.COV2.S vaccine regimes in Syrian hamsters. **a)** Syrian hamsters were immunized with either 10^9^ or 10^10^ VP (N=12 per dose level) of Ad26-based vaccines, or with 10^10^ vp of an Ad26 vector without gene insert as control (Ad26.empty, N=6). Four weeks after immunization half the hamsters per group received a second immunization with the same Ad26-based vaccine (N=6 per group). **b**) SARS-CoV-2 neutralization titers were measured 4 weeks after dose 1 and **c)** 4 weeks after dose 2 by wild-type VNA determining the inhibition of the cytopathic effect of SARS-CoV-2 on Vero E6 cells. The sera from Syrian hamsters immunized with Ad26.Empty were pooled into 2 groups for negative control samples. Median responses per group are indicated with horizontal lines. Dotted lines indicate the LLOD. Animals with a response at or below the LLOD are displayed as open symbols on the LLOD. CPE = cytopathic effect; LLOD = Lower Limit of Detection; p.i. = post inoculation; VNA = virus neutralization assay; VP = virus particles.

Binding antibodies measured by an enzyme-linked immunosorbent assay (ELISA) showed the same differences between Ad26.COV2.S and the two prototype vaccines (Supp Fig 2). However, a second dose at week 4 only transiently increased the median binding antibody titers at week 5. Antibody titers subsequently declined and at week 8 were comparable to levels observed prior to dose 2 at week 4 or lower.

We confirmed the immunogenicity of Ad26.COV2.S and the two prototype vaccines and the benefit of a second dose in rabbits (New Zealand white rabbits, female), in a 2-dose regimen using an 8-week interval, an interval that is also being evaluated in clinical studies^5^. Vaccines were tested at a dose level of, 5×10^9^ and 5×10^10^ VPs, of which the latter represents the human dose used in phase 3 clinical trials^5^. All tested vaccines elicited binding and neutralizing antibody titers as early as 2 weeks after the 1^st^ dose (no samples were collected earlier after immunization) with Ad26.COV2.S again inducing higher antibody titers compared to both Ad26.S and Ad26.dTM.PP (Supp Fig 3A and B). A second homologous dose at week 8 significantly boosted the binding and neutralizing antibody titers at week 10 (2 weeks post second dose) when compared across dose to pre-second dose levels (p<0.001).

**Figure 3.**
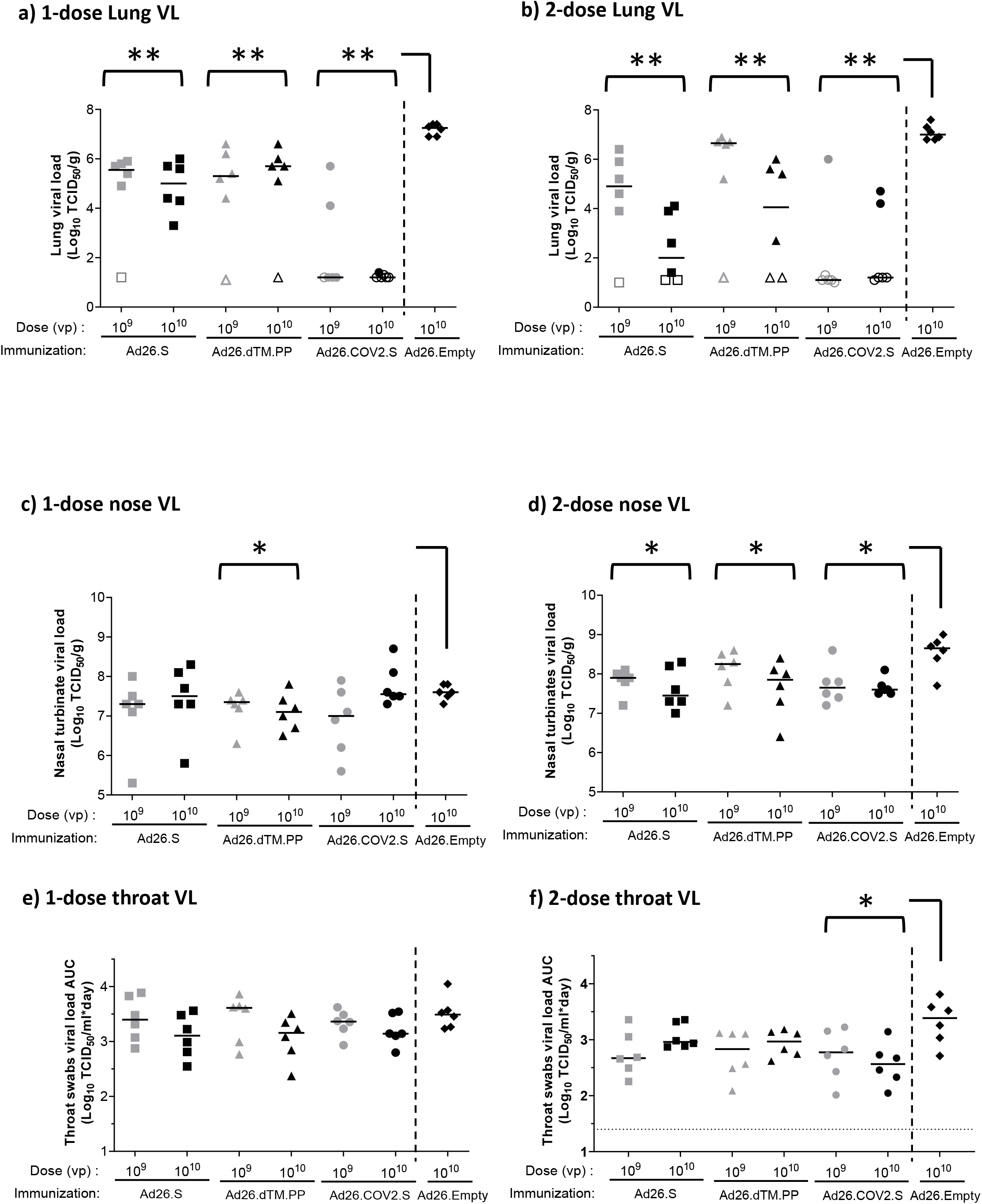
Protection against SARS-CoV-2 viral replication in Syrian hamsters immunized with Ad26-based vaccines. Syrian hamsters were intramuscularly immunized with a 1-dose regimen and a 2-dose regimen of Ad26.S, Ad26.dTM.PP, Ad26.COV2.S, or Ad26.empty (Ad26 vector not encoding any SARS-CoV-2 antigens). Hamsters received an intranasal inoculation with 10^2^ TCID_50_ SARS-CoV-2 strain BetaCoV/Munich/BavPat1/2020 4 weeks post-dose 1 (week 4) or 4 weeks post-dose 2 (week 8). **a, b)** Right lung tissue and **c, d)** right nasal turbinates were harvested at the end of the 4-day inoculation phase for viral load analysis. Replication competent virus was measured by TCID_50_ assay. **e, f)** Throat swab samples were taken daily after inoculation, and viral load area under the curve during the four-day follow-up was calculated as TCID_50_/ml x day. The median viral load per group is indicated with a horizontal line. LLOD was calculated per animal and animals with a response at or below the LLOD are shown as open symbols on the LLOD. Comparisons were performed between the Ad26.S, Ad26.dTM.PP and Ad26.COV2.S groups across dose level, with the Ad26.empty group by Mann-Whitney U-test. Statistical differences indicated by asterisks: *: p<0.05; **:p<0.01. LLOD = lower limit of detection; TCID_50_/g = 50% tissue culture infective dose per gram tissue; TCID_50_/ml = 50% tissue culture infective dose per ml sample; VP = virus particles.

### Protective efficacy of Ad26.COV2.S against SARS-CoV-2 challenge in Syrian hamsters

Next, we studied the protective efficacy of Ad26.COV2.S and the two prototype vaccines administered as 1- or 2-dose regimens followed by an intranasal inoculation with 10^2^ TCID_50_ of SARS-CoV-2 G614 virus 4 weeks after the last dose i.e., at week 4 for the 1-dose regiment and at week 8 for the 2-dose regimen in Syrian hamsters. At 4 days post inoculation (dpi), animals were sacrificed, and lungs, nasal turbinates and throat swabs were analyzed for viral load and pathological changes. To increase the power of the statistical analysis and since we did not observe a pronounced dose responsiveness for the virology readouts, we pooled viral load readouts for comparison to the Ad26.Empty control group. Comparisons between the three different vaccines (Ad26.COV2S, Ad26.S, Ad26.dTM.PP) were conducted across dose levels.

After a single vaccination and subsequent challenge, median lung viral load in all vaccine groups was significantly lower compared with the Ad26.Empty control group (median viral load 10^7.3^ TCID_50_/g) (Fig 3A). Virus was detected in the lungs of 11 out of 12 animals immunized with a single dose of Ad26.S (median viral load 10^9^ VP 10^5.6^ TCID_50_/g, 10^10^ VP 10^5^ TCID_50_/g), and in 10 out of 12 animals immunized with a single dose of A26.dTM.PP (median viral load 10^9^ VP 10^5.3^ TCID_50_/g, 10^10^ VP 10^5.7^ TCID_50_/g) and no clear effect of vaccine dose was observed. By contrast, only 3 out of 12 animals immunized with a single dose of Ad26.COV2.S had detectable virus in the lung (median viral load of 10^9^ VP and 10^10^ VP 10^1.2^ TCID_50_/g). Of hamsters vaccinated with a second dose of Ad26.COV2.S, again 3 out of 12 animals had detectable viral load for both vaccine dose levels, suggesting no added value of the second dose. Viral load in the animals with breakthrough infections were also similar to the viral load in animals with breakthrough infections after one vaccination. A second dose of 10^9^ VP Ad26.S or Ad26.dTM.PP also had little impact on the lung infection rate post challenge at week 8 (5 out of 6 animals showed detectable viral load per vaccine, median titer 10^4.9^ TCID_50_/g and 10^6.7^ TCID_50_/g, respectively) but a second dose of 10^10^ VP of these prototype vaccines was associated with a lower lung infection rate and median lung viral loads compared with the 1-dose regimens (median titer 10^2^ TCID_50_/g and 10^4^ TCID_50_/g, respectively), suggesting a benefit of a 2 dose regimen (Fig 3B).

To determine the impact of the vaccines on viral load in the upper respiratory tract, nasal turbinate viral load was determined after sacrifice at 4 dpi, and throat swab viral load was determined daily after infection and was analyzed as area under the curve (AUC) per animal up to day 4 post infection. In animals receiving a single vaccine dose, a limited but statistically significant reduction in nasal turbinate viral load after challenge was observed for Ad26.dTM.PP but not for Ad26.COV2.S and Ad26.S compared with the control group. After 2 vaccine doses, all three vaccines induced a significant reduction in nasal turbinate viral load post challenge compared with the Ad26.Empty group (Fig 3C and D). By contrast, throat swab viral load data show that none of the vaccines reduced viral burden in the throat after single immunization and subsequent inoculation with SARS-CoV-2 (Fig 3E), and only animals immunized with two doses Ad26.COV2.S had significantly reduced throat viral load compared to control (Fig 3F).

The observed protective efficacy results are further supported by immunohistochemistry (IHC) staining for SARS-CoV-2 NP in the lung and nose tissue, and by histopathology (Supp Fig 4 and 5). Lung and nose IHC and histopathology scores were overall consistent with viral load data, with lower median scores in the lungs of immunized groups compared with the Ad26.Emtpy group (Supp Fig 4, table 1) and no significant difference in the IHC and histopathology scores in nose tissues of vaccinated animals compared with the Ad26.empty control group independent of vaccine regimen (Supp Fig 5, table 1).

### Syrian hamsters immunized with sub-optimal dose levels Ad26.COV2.S do not show signs of VAERD after SARS-CoV-2 inoculation and breakthrough infection

Based on the observed immunogenicity and efficacy, Ad26.COV2.S was selected for further evaluation in a dose titration study to address the theoretical risk of VAERD under conditions of suboptimal immune responses allowing breakthrough infection after SARS-CoV-2 challenge. Groups of hamsters were immunized with a single dose of Ad26.COV2.S at 10^7^, 10^8^, 10^9^ or 10^10^ VP (4 groups; n=8/group). The control group received 10^10^ VP of a control vaccine encoding an irrelevant antigen (Ad26.Irr). At 4 weeks post immunization we observed a clear dose response of binding and neutralizing antibodies, both in number of responding animals and in antibody titers, with no detectable binding or neutralizing antibody titers in 1 or 3 out of 8 animals at the lowest dose level of 10^7^ VP (Fig 4A and B), respectively. For the 10^9^ and 10^10^ VP doses, median levels of binding antibodies (median endpoint titer of 10^3.7^ and 10^3.9^, respectively) and median neutralizing antibody responses (median neutralization titers of 45 and 91, respectively) were consistent with observations in the previous study. Four weeks after immunization, hamsters were inoculated with 10^2^ TCID50 SARS-CoV-2 followed by determination of efficacy and histopathology readouts as in the previous study. Animals dosed with 10^9^ and 10^10^ VP of Ad26.COV2.S showed similar frequencies of breakthrough lung infection as the comparable groups in the previous study (Fig 3A), with 3 out of 8 and 2 out of 8 animals with detectable lung viral load in the dose titration study, respectively (Fig 4C). Despite the increase in the number of animals that had breakthrough infections at lower Ad26.COV2.S dose levels (6 out of 8 animals that received 10^8^ VP and 8 out of 8 that received 10^7^ VP), the median lung viral load titers (median of 10^4.8^ TCID_50_/g at 10^8^ VP, median of 10^5^ TCID_50_/g at 10^7^ VP) and IHC staining of SARS-CoV-2 NP in these groups (median scores 1) were lower than in the control group (median lung viral load 10^6.8^ TCID_50_/g and IHC median score 2) (Fig 4C and D). Congruent with lung viral load and IHC staining results, immunization with 10^8^, 10^9^ and 10^10^ VP significantly reduced histopathology in the lower respiratory tract compared with mock-immunized hamsters (Fig 5A). Immunization with 10^9^ and 10^10^ VP resulted in absence of any signs of lower respiratory tract histopathology in 4 out of 8 and 3 out of 8 hamsters, respectively. Notably, despite detectable breakthrough lung infection in all hamsters dosed with 10^7^ VP and in most hamsters immunized with 10^8^ VP, median lower respiratory tract histopathology scores were lower when compared with the mock immunized group (Ad26.irr, Table 2).

**Figure 4.**
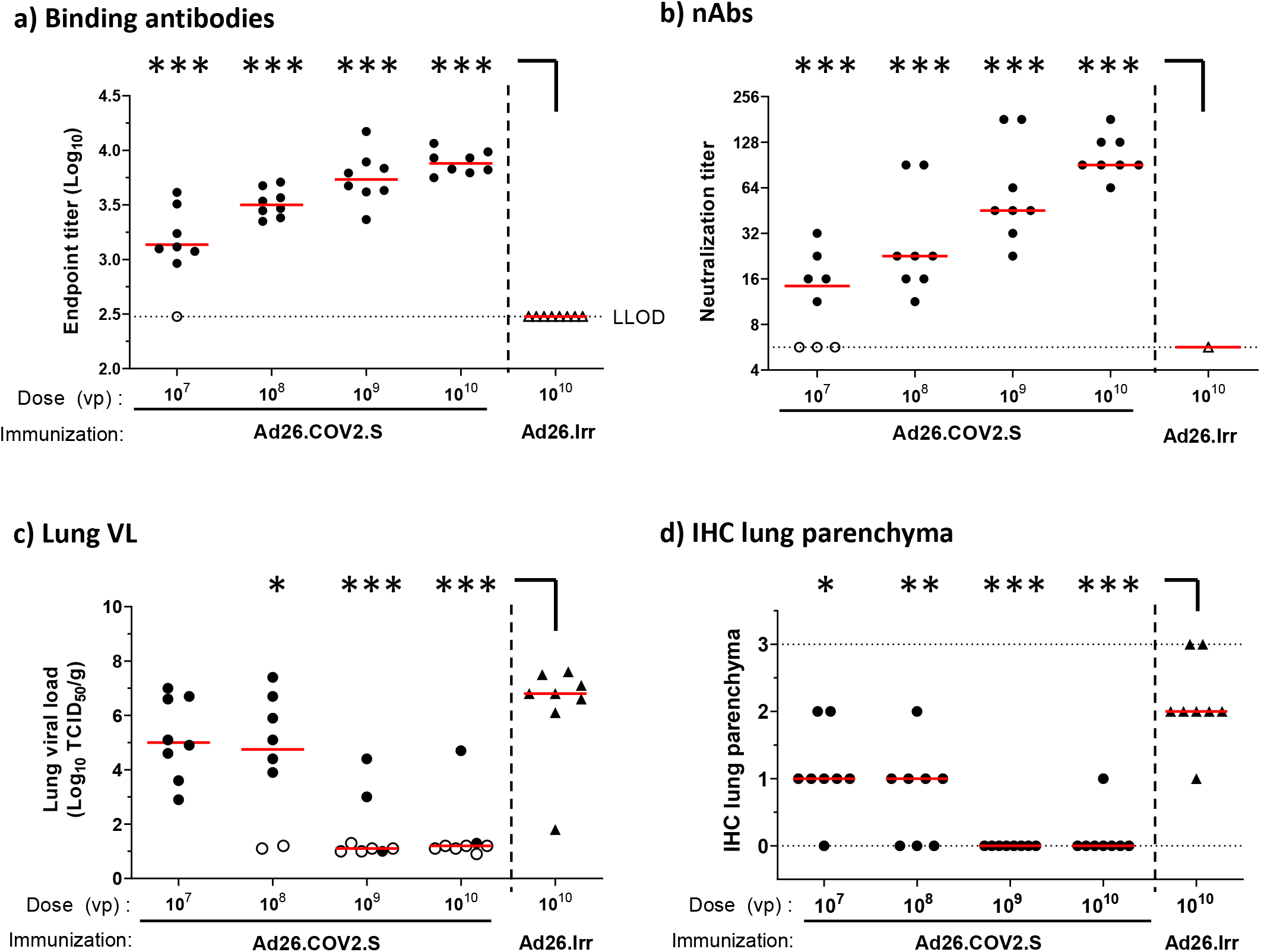
Dose responsiveness of Ad26.COV2.S on immunogenicity and lung viral load in hamsters. Syrian hamsters were intramuscularly immunized with 10^7^, 10^8^, 10^9^ or 10^10^ VP of Ad26.COV2.S N=8 per group, or 10^10^ VP Ad26.Irr (an Ad26 vector not encoding any SARS-CoV-2 antigens, N=8). Four weeks after one immunization, SARS-CoV-2 Spike protein-specific antibody binding titers as measured by ELISA (**a**) and SARS-CoV-2 neutralizing antibodies as measured by wtVNA (**b**) were determined. The median antibody responses per group is indicated with a horizontal line. Dotted lines indicate the LLOD. Animals with a response at or below the LLOD were put on LLOD and are shown as open symbols. Hamsters received intranasal inoculation with 10^2^ TCID_50_ SARS-CoV-2 strain BetaCoV/Munich/BavPat1/2020 4 weeks post immunization (week 4). Right lung tissue was isolated 4 days after inoculation for virological analysis and immunohistochemistry. **c)** Lung viral load was determined by TCID_50_ assay on Vero E6 cells. The median viral load per group is indicated with a horizontal line. LLOD was calculated per animal, and animals with a response at or below the LLOD are shown as open symbols. **d)** presence of SARS-CoV-2 NP was determined by immunohistochemical staining. Comparisons were performed between the Ad26.COV2.S dose level groups, with the Ad26.Irr group by Mann-Whitney U-test. Statistical differences indicated by asterisks: *:p<0.05; **:p<0.01; ***:p<0.001. Ad26.Irr = Ad26 vector not encoding any SARS-CoV-2 antigens; LLOD = lower limit of detection; N = number of animals; TCID_50_/g = 50% tissue culture infective dose per gram tissue; VP = virus particles; NP = Nucleocapsid protein; wtVNA = wild-type virus neutralization assay.

**Figure 5.**
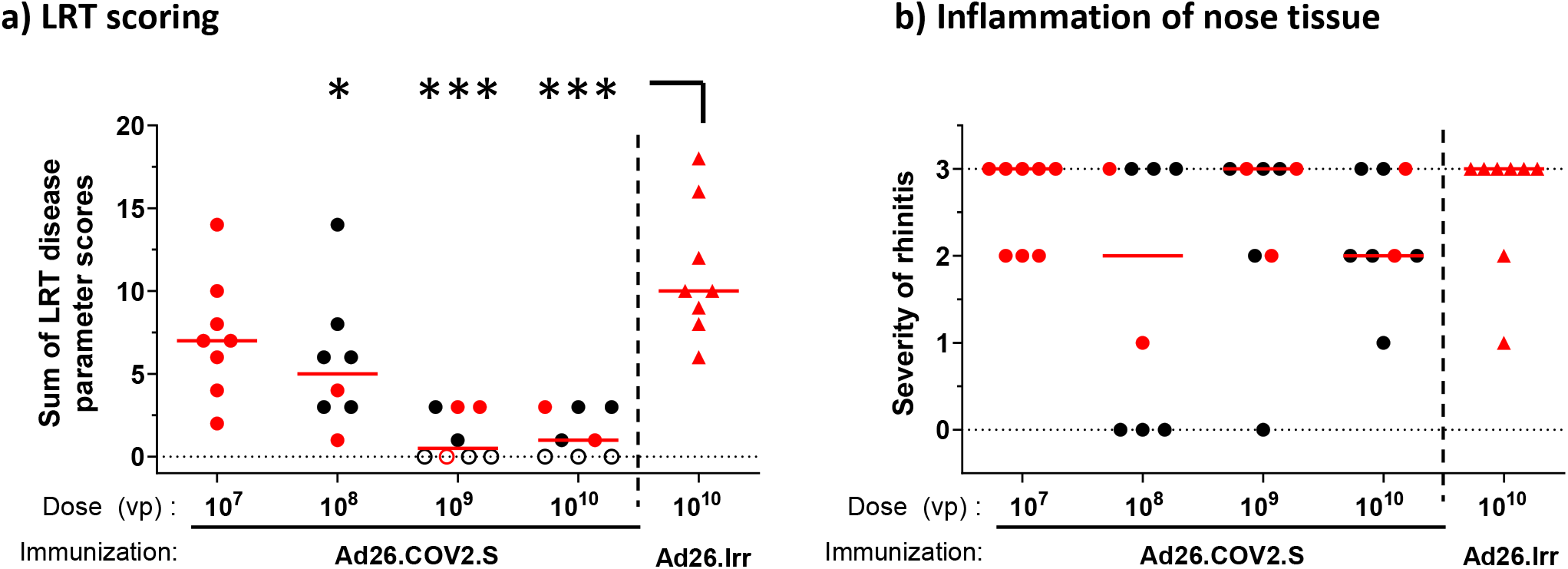
No signs of VAERD in Ad26 immunized Syrian hamsters inoculated with SARS-CoV-2. Four days after IN inoculation with 10^2^ TCID50 SARS-CoV-2 (N = 8 per group), **a)** lung tissue was isolated and scored for presence and severity of alveolitis, alveolar damage, alveolar edema, alveolar hemorrhage, type II pneumocyte hyperplasia, bronchitis, bronchiolitis, peribronchial and perivascular cuffing. Sum of scores are presented as sum of LRT disease parameters. **b)** Four days after inoculation, nose tissue was isolated and scored for severity of inflammation (rhinitis). Horizontal lines denote a pathology score of 0, indicating no histopathology. Symbols in red denote samples from hamsters with breakthrough lung viral load (>10^2^ TCID_50_/g). Comparisons were performed between the Ad26.COV2.S dose level groups, with the Ad26.Irr group by Mann-Whitney U-test. Statistical differences indicated by asterisks: *:p<0.05; **:p<0.01; ***:p<0.001. Ad26.Irr = Ad26 vector not encoding any SARS-CoV-2 antigens; LRT = lower respiratory tract; N = number of animals; VP = virus particles.

The inflammation score of nasal tissue (rhinitis) showed no significant differences between vaccinated and control groups (Fig 5B). Collectively, these data demonstrate that the presence of low levels of neutralizing antibodies elicited by sub-optimal Ad26.COV2.S vaccine dose levels do not aggravate lung disease in challenged Syrian hamsters when compared to a mock vaccine.

### Binding and neutralizing antibodies correlate with protection

To determine putative correlates of protection, binding and neutralizing antibody titers from different regimens and dose levels were pooled for Ad26.COV2.S (N=56) and compared between protected and unprotected animals (Fig 6). Protection from SARS-CoV-2 infection was defined as a lung viral load below 10^2^ TCID/g, based on the observation that only few animals with detectable viral load fall below this margin, which was likely related to variation in the available sample quantity per animal (Fig 3A and B, and Fig 4C). Protected animals dosed with Ad26.COV2.S had significantly (2.3-fold) higher median binding antibody titers than unprotected animals (p<0.001, two sample t-test) (Fig 6A). Similar results were observed for an analogous analysis of median neutralizing antibody titers, which were also significantly (4-fold) increased in animals immunized with Ad26.COV2.S with undetectable lung viral load compared with unprotected animals (p<0.001, two sample t-test) (Fig 6B).

**Figure 6.**
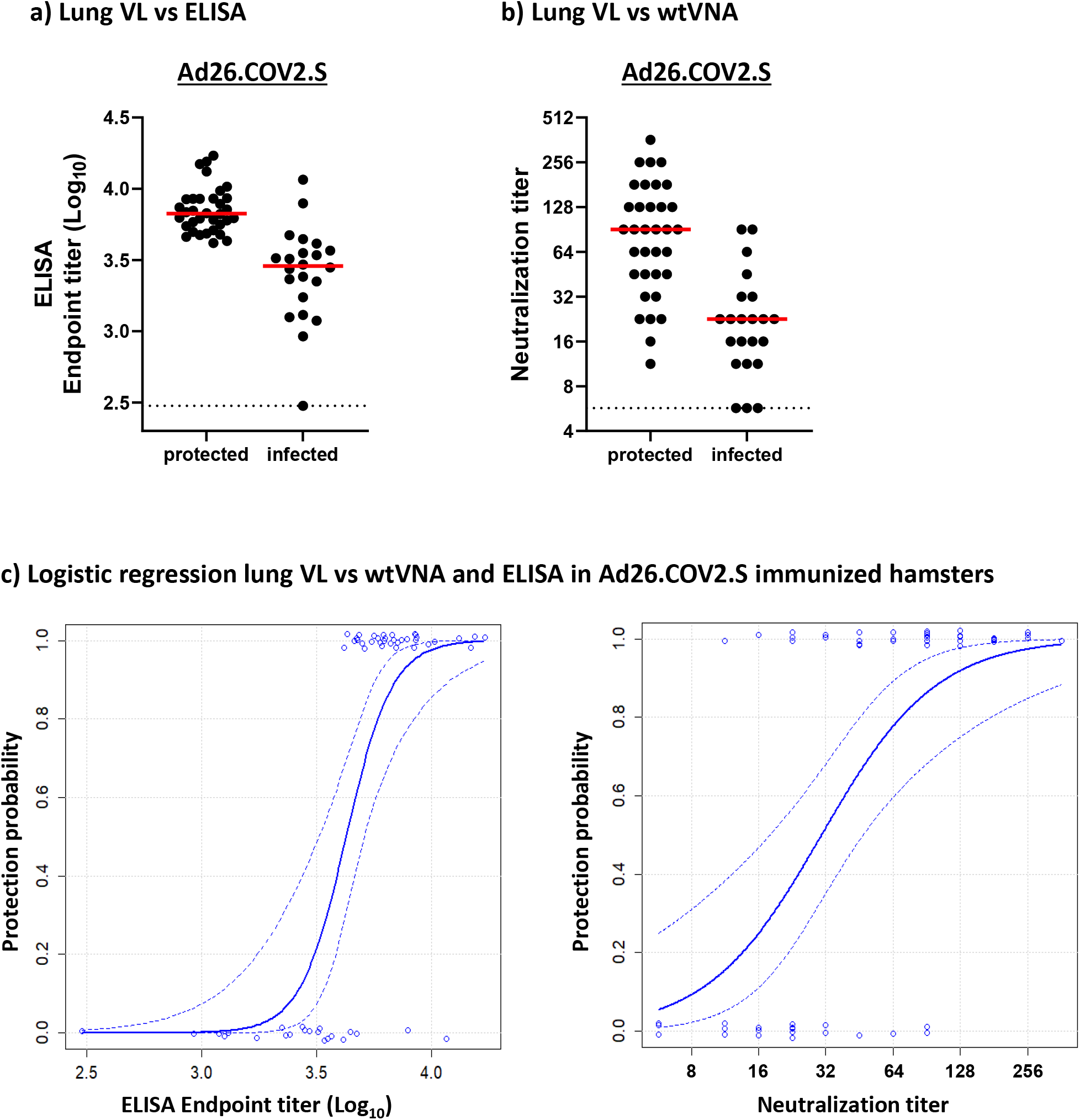
Binding and neutralizing antibodies correlate with protection. Protection was defined as a viral load below 10^2^ TCID_50_/g in lung tissue, irrespective of vaccine regimen and dose level (see Fig 3A and B, and Fig 4C). Syrian hamsters were immunized once, or twice, with 10^7^, 10^8^, 10^9^, 10^10^ VP Ad26.CoV2.S (N=56). Hamsters were inoculated with 10^2^ TCID_50_ SARS-CoV-2, and four days later sacrificed for virological analysis of lung tissue. Prior to virus inoculation serum samples were analyzed for **a)** antibody binding titers and **b)** virus neutralizing antibodies. Median antibody responses per group is indicated with horizontal lines. Dotted lines indicate the LLOD. **c)**) Logistic regression models using Firth’s correction were built with protection outcome as the dependent variable, and binding and neutralizing antibody titers from pooled regimens and dose levels of Ad26.COV2.S as independent variable. Dotted lines indicate the 95% confidence interval. LLOD = lower limit of detection; N = number of animals; TCID50/g = 50% tissue culture infective dose per gram tissue; VP = virus particles.

To gain a more quantitative understanding of the relationship between immune response levels and protection outcome, we built logistic regression models with Firth’s correction (Fig 6C). Hamsters were classified either as infected or protected from SARS-CoV-2, as defined above. Binding antibody titers correlated significantly with protection (p=0.0004), with endpoint titers above 10^3.6^ appearing to be linked with protection from lung infection. Also a comparably significant slope (p=0.0002) was observed with neutralizing antibody titers where titers above 32 appeared to be linked with protection outcome in the majority of animals.

## DISCUSSION

In a previous study using an early SARS-CoV-2 isolate we have demonstrated that immune responses elicited by a single dose of Ad26.COV2.S could reduce viral load and protected hamsters from severe clinical disease^3^. However, during the ongoing SARS-CoV-2 pandemic, a virus variant with a D614G substitution in the spike protein has emerged^16^. This mutation has been associated with increased viral fitness and enhanced infectivity and has now become the dominant variant in large parts of the world^16^, although likely to be replaced over time by new variants that are constantly emerging. In Syrian hamsters, it was confirmed that infection with the D614G variant was associated with higher infectious viral titers in the upper respiratory tract but not in the lungs ^21^. Here we describe the establishment of a hamster challenge model of moderate disease using a SARS-CoV-2 strain containing the prevalent D614G substitution. We used this model to test the protective efficacy of immune responses elicited by our COVID-19 vaccine candidate Ad26.COV2.S and two Ad26-based prototype vaccines that encode different SARS-CoV-2 spike designs. Our study demonstrates that the disease progression in this hamster challenge model shows features of a moderate disease course in humans with clear histopathological lung disease which was only marginally exacerbated by a larger inoculum dose. Peak lung viral load was not affected by a lower inoculum dose, suggesting infection of the majority of primary susceptible lung cells, leading to peak lung viral load at day 2 post inoculation. In line with our previous studies in vaccinated and challenged NHPs and hamsters, Ad26.COV2.S vaccination reduced viral replication in lungs by 6 log_10_ below the level observed in control animals with many animals that received higher vaccine doses showing undetectable viral replication. Ad26.COV2.S significantly outperformed the two prototype vaccines, both for immunogenicity as for protective efficacy. Our published data^3,37^ in combination with our present study indicate that Ad26.COV2.S-elicited immune responses give adequate protection against SARS-CoV-2 variants with and without the D614G spike substitution.

We extended previous studies by evaluation of a second homologous vaccine dose with a 4 week interval, which only moderately increased binding antibody levels, while neutralizing antibody titers were substantially boosted. This contrasts observations in NHP, where a second vaccine dose significantly boosted both neutralizing and binding antibody levels^38^. Possible explanations include limited translatability of dose levels between hamsters and NHP, differential impact of anti-Ad26 vector responses elicited by the first dose on the immunogenicity of a second homologous dose between species, and that a shorter interval of 4 weeks between immunizations compared with 8 weeks can reduce the impact of a second dose, as previously observed in NHP^37^. In addition, the high binding antibody levels induced by a single immunization in hamsters might represent saturating levels while neutralizing antibodies could still increase after a second dose, possibly reflecting extended affinity maturation. The advantage of a second homologous vaccine dose for humoral SARS-CoV-2 S-specific immune responses was also observed in rabbits immunized with the same Ad26-based vaccines and confirm our clinical data (Sadoff, le Gars, NEJM in press). Whether a 2-dose regimen is also preferred for improved vaccine efficacy remains to be seen.

Interestingly, a second dose of Ad26.S or Ad26.dTM.PP increased protection against lung viral load after challenge compared with the low protection achieved by a single vaccination of hamsters with these prototype vaccines. In contrast, a second dose of Ad26.COV2.S did not further increase the already high level of efficacy established by a single dose. This is supported by our correlate analysis where the probability of protection increases with a higher antibody titer, and if a certain antibody titer is reached, protection probability increases only moderately. The correlation of lung protection with serum binding- and neutralizing antibody levels, as observed here in the Syrian hamster SARS-CoV-2-D614G challenge study, confirms our data in NHP^2^ and was irrespective of vaccine, dose level or regimen.

In addition to protection from COVID-19, vaccine-elicited immunity ideally also protects against asymptomatic infection as well as against transmission of virus by reduction of viral load in the upper respiratory tract. In the hamster challenge model used here we observed high replication-competent virus levels in the nasal turbinates despite the low virus inoculum dose used, which is in line with the observation that the spike D614G substitution increases SARS-CoV-2 infectivity in the upper respiratory tract of challenged hamsters^21^. As the size of the challenge inoculum is low it is unlikely that the high viral load in nasal turbinates detected 4 days later are derived from the original inoculum. Viral load reduction in nose tissue required two vaccine doses, irrespective of the vaccine used. Two doses of Ad26.COV2.S was the only regimen that also decreased viral titers in throat swabs. Reduction of viral load in the upper respiratory tract was limited compared to the lower respiratory tract which is in contrast to our NHP studies where we observed almost complete reduction of nasal viral load. This may be explained by a difference in the susceptibility of the nasal epithelium for viral infection^41,42^, or the potentially different composition of immune cells present in the respiratory tract between hamsters and primates and different induction of local upper respiratory tract immunity by vaccine candidates in general or by Ad26-based vaccines specifically. Whether Ad26.COV2.S-elicited immunity can protect against asymptomatic infection and SARS-CoV-2 transmission remains to be determined.

Previous studies in preclinical models with candidate coronavirus vaccines against SARS-CoV and MERS indicated that disease can be exacerbated upon infection by certain vaccine-elicited immune responses^43^. However, neither VAERD nor antibody-mediated disease enhancement have been reported following vaccination with SARS-CoV-2 vaccine candidates in pre-clinical animal models, nor in ongoing clinical studies including efficacy reports of phase 3 studies of mRNA- and other adenoviral vector-based or whole inactivated virus vaccines. Nevertheless, vaccine efficacy in clinical studies was high so far and the theoretical potential for VAERD requires further investigation especially in the setting of suboptimal or waning vaccine-induced immunity. We therefore assessed the potential for Ad26.COV2.S to predispose for VAERD in a setting where levels of vaccine-induced antibodies were too low to prevent viral replication in the lung by immunizing with suboptimal Ad26.COV2.S doses. Importantly, even in the setting of inadequate immune responses for the prevention of lung viral replication, the lower respiratory tract histopathology scores of immunized animals showed no signs of VAERD when compared to the control group. Conversely, most vaccinated animals with breakthrough infection still showed reduced histopathology compared with control animals. These results imply that the theoretical risk that Ad26.COV2.S would predispose for VAERD is minimal.

Our study confirms that our Ad26.COV2.S vaccine candidate is highly immunogenic, and can protect hamsters against challenge with a SARS-CoV-2 G614 spike variant virus. The excellent potency of Ad26.COV2.S and the absence of data that it would predispose for VAERD, supports its continuous evaluation in the ongoing Phase 3 clinical trials in a single and a two-dose regimen (NCT04505722 and NCT04614948, respectively).

## MATERIALS AND METHODS

### Vaccines

The Ad26-based vaccines were generated as previously described^4^. Briefly, they are based on a replication incompetent adenovirus serotype 26 (Ad26) vector encoding a prefusion stabilized SARS-COV-2 spike protein sequence (Wuhan Hu1; GenBank accession number: MN908947). Replication-incompetent, E1/E3-deleted Ad26-vectors were engineered using the AdVac system^44^, using a single plasmid technology containing the Ad26 vector genome including a transgene expression cassette. The codon optimized, prefusion stabilized, SARS-COV-2 spike protein encoding gene was inserted into the E1-position of the Ad26 vector genome. Manufacturing of the Ad26 vectors was performed in the complementing cell line PER.C6 TetR ^45,46^. The negative control vector Ad26.Irr (RSV-FA2-2A-GLuc) encodes the RSV F protein fused to Gaussia firefly luciferase as a single transgene separated by a 2A peptide sequence, resulting in expression of both individual proteins. Manufacturing of the vector was performed in Per.C6. Adenoviral vectors were tested for bioburden and endotoxin levels prior to use.

### Study design animal experiments

#### Hamster studies

Animal experiments were approved by the Central Authority for Scientific Procedures on Animals (Centrale Commissie Dierproeven) and conducted in accordance with the European guidelines (EU directive on animal testing 86/609/EEC) and local Dutch legislation on animal experiments. The in-life phase took place at Viroclinics Biosciences BV, Viroclinics Xplore, Schaijk, the Netherlands. All Viroclinics personnel involved in performing the clinical observations and laboratory analysis in which interpretation of the data was required were not aware of the Treatment Allocation Key at any time prior to completion of the study and were blinded by allocating a unique sample number to each sample collected and analysis.

Male Syrian (golden) hamsters (*Mesocricetus auratus*), strain HsdHan:AURA, aged 9-11 weeks at the start of the study were purchased from Envigo (Envigo RMS B.V., Venray, the Netherland). Hamsters were immunized via the intramuscular route with 100μl vaccine (50μl per hind leg) under isoflurane anesthesia. Hamsters were intranasally inoculated with 100μl containing 10^2^ TCID_50_ of SARS-CoV-2 (BetaCoV/Munich/BavPat1/2020, containing a D614G substitution in the S1 fragment, kindly provided by Dr. C. Drosten). The sequence of the challenge stock has been characterized and has been shown to be in line with the parental strain (data not shown). On the day of infection, prior to inoculation, and daily until four days post infection throat swabs were collected under isoflurane anesthesia. Throat swabs were collected in virus transport medium, aliquoted and stored until time of analysis. Intermediate blood samples were collected via the retro-orbital bleeding route under isoflurane anesthesia. Blood was processed for serum isolation. At the end of the experiment, under anesthesia, animals were sacrificed by cervical dislocation and necropsy was performed. Respiratory tissues collected after necropsy were analyzed for viral load, and for histopathological changes.

#### Rabbit studies

Rabbit experiments were approved by the local animal welfare body and conducted in concordance with European guidelines (EU directive on the protection of animals used for scientific purposes 2010/63/EU) and local Belgian legislation on animal experiments. The in-life phase took place at the non-clinical safety Beerse site of Janssen Research and Development, an AAALAC-approved laboratory. Female New Zealand White rabbits, aged approximately 4 months at the start of the study were purchased from Charles River Laboratories in France. Rabbits were immunized in week 0 and week 8 of the study with 5×10^9^ or 5×10^10^ vp Ad26.S, Ad26.dTM.PP or Ad26.COV2.S in a volume of 0.5 mL via the intramuscular route. As a control group, five rabbits were immunized with saline. Interim blood samples for serum processing were collected via the lateral vein in the ear. At the end of the experiment, animals were sacrificed by intravenous injection of Sodiumpentobarbital, followed by exsanguination via the femoral artery.

### Detection of infectious viral load by TCID_50_ assay

Quadruplicate 10-fold serial dilutions were used to determine the TCID_50_ virus titers in confluent layers of Vero E6 cells. To this end, serial dilutions of the samples (throat swabs, and tissue homogenates) were made and incubated on Vero E6 monolayers for 1 hour at 37 °C. Vero E6 monolayers are washed and incubated for 5-6 days at 37 degrees after which plates are scored using the vitality marker WST-8 (colorimetric cell counting kit, Sigma Aldrich, cat 96992-3000TESTS-F). To this end, WST-8 stock solution was prepared and added to the plates. Per well, 20 μL of this solution (containing 4 μL of the ready-to-use WST-8 solution from the kit and 16 μL infection medium, 1:5 dilution) was added and incubated 3-5 hours at room temperature. Subsequently, plates were measured for optical density at 450 nm (OD450) using a micro plate reader and visual results of the positive controls (cytopathic effect (cpe)) were used to set the limits of the WST-8 staining (OD value associated with cpe). Viral titers (TCID_50_) were calculated using the method of Spearman-Karber.

### Histopathology

Histopathology was assessed by a pathologist from Viroclinics Biosciences BV, Viroclinics Xplore, and a pathologist from Janssen Non-Clinical Safety (Beerse, Belgium).

Four days p.i. all animals were autopsied by opening the thoracic and abdominal cavities and examining all major organs. The extent of pulmonary consolidation was assessed based on visual estimation of the percentage of affected lung tissue. The left nasal turbinates, trachea and left lung were collected for histopathological examination and analysis by IHC. All tissues were gently instilled with, and/or immersed in 10% neutral-buffered formalin for fixation. Lungs and trachea were routinely processed, paraffin wax embedded, micro-sectioned to 3 μm on glass slides and stained with haematoxylin and eosin (H&E) for histopathological evaluation. The sampled and fixed nasal turbinates were processed after decalcification and embedded into paraffin blocks, and similarly cut and stained. The H&E stained tissue sections were examined by light microscopy for histopathology scoring, as well as for the presence of any other lesions. The severity of inflammatory cell infiltration in nasal turbinates and tracheas was scored for rhinitis and tracheitis: 0 = no inflammatory cells, 1 = few inflammatory cells, 2 = moderate number of inflammatory cells, 3 = many inflammatory cells.

For lung tissue, each entire slide was examined and scored for presence or absence of alveolar edema, alveolar hemorrhage and type II pneumocyte hyperplasia (0 = no, 1 = yes). The degree and severity of inflammatory cell infiltration and damage in alveoli, bronchi/bronchioles were scored for alveolitis and bronchitis/bronchiolitis: 0 = no inflammatory cells, 1 = few inflammatory cells, 2 = moderate number of inflammatory cells, 3 = many inflammatory cells. Extent of peribronchial/perivascular cuffing: 0 = none, 1 = 1-2 cells thick, 2 = 3-10 cells thick, 3 = over 10 cells thick. Additionally, the extent of alveolitis/alveolar damage was scored per slide: 0 = 0%, 1 = <25%, 2 = 25-50%, 3 = >50%.

The cumulative score (sum) for the extent and severity of inflammation of lung tissues provided the total lower respiratory tract (LRT) score, with a possible maximum score of 24. The following histopathology parameters were included in the sum of lower respiratory tract disease parameters: alveolitis, alveolar damage, alveolar edema, alveolar hemorrhage, type II pneumocyte hyperplasia, bronchitis, bronchiolitis, peribronchial and perivascular cuffing.

### Immunohistochemistry

Lung, nose and trachea tissue samples were sampled, fixed in 10% formalin (lung instilled) for 14 days and were embedded in paraffin by Viroclinics Biosciences B.V. Tissue blocks were delivered and assessed by a pathologist from Janssen Non-clinical Safety (Beerse, Belgium).

Paraffin sections of lung, trachea and nose sections of all animals were automatically stained (Ventana Discovery Ultra, Roche, France), using rabbit polyclonal anti-SARS-CoV Nucleocapsid protein antibody (NP, Novus NB100-56576, 1/300) which is cross reactive towards SARS-CoV-2 NP. These sections were semi-quantitatively scored for number of SARS-CoV-2 NP positive cells, and graded as 0: no positive immunoreactive cells, 1: minimal (few/focal) number of positive cells, 2 moderate (focal/multifocal) number of positive cells and 3: many/high (focally extensive/multifocal) number of immunoreactive cells.

### Virus Neutralization Assay

Neutralization assays against live SARS-CoV-2 were performed using the microneutralization assay previously described by Algaissi and Hashem^47^. Vero E6 cells [CRL-1580, American Type Culture Collection (ATCC)] were grown in Eagle’s minimal essential medium (EMEM; Lonza) supplemented with 8% fetal calf serum (FCS; Bodinco BV), 1% penicillin-streptomycin (Sigma-Aldrich, P4458) and 2 mM L-glutamine (PAA). Cells were maintained at 37°C in a humidified atmosphere containing 5% CO2. Clinical isolate SARS-CoV-2/human/NLD/Leiden-0008/2020 (Leiden L-0008) was isolated from a nasopharyngeal sample and its characterization will be described elsewhere (manuscript in preparation). Isolate Leiden-0008 was propagated and titrated in Vero E6 cells using the TCID_50_ endpoint dilution method. The next-generation sequencing derived sequence of this virus isolate is available under GenBank accession number MT705206 and shows 1 mutation in the Leiden-0008 virus spike protein compared to the Wuhan spike protein sequence resulting in Asp>Gly at position 614 (D614G) of the Spike protein. In addition, several non-silent (C12846U and C18928U) and silent mutations (C241U, C3037U and C1448U) in other genes were found. The TCID_50_ was calculated by the Spearman-Kärber algorithm as described^48^. All work with live SARS-CoV-2 was performed in a biosafety level 3 facility at Leiden University Medical Center.

Vero-E6 cells were seeded at 12,000 cells/well in 96-well tissue culture plates 1 day prior to infection. Heat-inactivated (30 min at 56°C) serum samples were analyzed in duplicate. The panel of sera were 2-fold serially diluted in duplicate, with an initial dilution of 1:10 and a final dilution of 1:1280 in 60 μL EMEM medium supplemented with penicillin, streptomycin, 2 mM L-glutamine and 2% FCS. Diluted sera were mixed with equal volumes of 120 TCID_50_/60 μL Leiden −0008 virus and incubated for 1 h at 37 °C. The virus-serum mixtures were then added onto Vero-E6 cell monolayers and incubated at 37 °C. Cells either unexposed to the virus or mixed with 120 TCID_50_/60 μL SARS-CoV-2 were used as negative (uninfected) and positive (infected) controls, respectively. At 3 days post-infection, cells were fixed and inactivated with 40 μL 37% formaldehyde/PBS solution/well overnight at 4 °C. The fixative was removed from cells and the clusters were stained with 50 μL/well crystal violet solution, incubated for 10 minutes and rinsed with water. Dried plates were evaluated for viral cytopathic effect. Neutralization titer was calculated by dividing the number of positive wells with complete inhibition of the virus-induced cytopathogenic effect, by the number of replicates, and adding 2.5 to stabilize the calculated ratio. The neutralizing antibody titer was defined as the log2 reciprocal of this value. A SARS-CoV-2 back-titration was included with each assay run to confirm that the dose of the used inoculum was within the acceptable range of 30 to 300 TCID_50_.

### ELISA

IgG binding to SARS-CoV-2 Spike antigen was measured by ELISA with an in-house produced COR200099 and COR200153 are SARS-CoV-2 spike proteins based on the Wuhan-Hu-1 SARS-CoV-2 strain (MN908947) and stabilized by two point mutations (R682A, R685G) in the S1/S2 junction that knock out the furin cleavage site, and by two introduced prolines (K986P, V987P) in the hinge region in S2. In addition, the transmembrane and cytoplasmic regions have been replaced by a foldon domain for trimerization, allowing the proteins to be produced as soluble proteins. COR200153 additionally contains an A942P mutation, which increases trimer expression and a C-terminal biotin label, which was covalently attached via a sortase A reaction.

For the analysis of hamster samples, 96-wells Perkin Elmer white ½ area plates were coated overnight with protein. For the analysis of rabbit samples, plates were incubated for 2 hours at 37°C for coating. Following incubation, plates were washed, blocked for 1 hour and subsequently incubated for 1 hour with 3-fold serially diluted serum samples in block buffer in duplicate. After washing, plates were incubated for 1 hour with Rabbit-Anti-Hamster IgG HRP (Invitrogen, catalogue number A18895) or anti-rabbit IgG-HRP (Jackson ImmunoResearch) in block buffer, washed again and developed using ECL substrate. Luminescence readout was performed using a BioTek Synergy Neo instrument (hamster samples) or on an Envision Multimode plate reader (rabbit samples). Hamster antibody titers are reported as Log_10_ endpoint, rabbit titers are reported as Log_10_ relative potency compared to a reference standard.

### Statistical analysis

Statistical differences across dose levels between immunization regimens were evaluated two-sided for S-specific binding antibodies as measured by ELISA, neutralizing titers as measured by virus neutralization assay (VNA), viral load as measured by TCID_50_, histopathology and IHC scores. Across dose levels comparisons between Ad26.S, Ad26.dTM.PP and Ad26.COV2.S groups were made using the t-test from ANOVA with vaccine and dose as factors for group comparisons without censored measurements at LLOD or LLOQ, or the z-test from Tobit ANOVA for group comparisons with at most 50% censored values, or the Cochran-Mantel-Haenszel test for group comparisons with 50% or more censored values. Results were corrected for multiple comparisons by 3-fold Bonferroni correction. Exploratory comparisons per dose level, and across dose level of Ad26.S, Ad26.dTM.PP and Ad26.COV2.S groups with groups immunized with an irrelevant antigen, Ad26.Empty and Ad26.Irr, were made using the methods above or the Mann-Whitney U test. Due to the exploratory nature of these analysis, results were not corrected for multiple comparisons.

Statistical analyses were performed using SAS version 9.4 (SAS Institute Inc. Cary, NC, US) and R version 3.6.1 (2019-07-05). Statistical tests were conducted two-sided at an overall significance level of α = 0.05.

### Correlation analysis

Hamsters were classified either as infected or protected from SARS-CoV-2, defined as a lung viral load of either above or below 10^2^ TCID_50_/g, respectively. From the binding and neutralizing antibody data pooled from different regimens and dose levels of Ad26.COV2.S, logistic regression models were built with Firth’s correction^49^, with protection outcome as the dependent variable, and the wtVNA and Log_10_ transformed ELISA data before inoculation as the independent variable.

## Acknowledgements

This project was funded in part by the Department of Health and Human Services Biomedical Advanced Research and Development Authority (BARDA) under contract HHS0100201700018C.

We thank Susan King, Joanna Bleszynska, Sanne Kroos, Sven Blokland, Ava Sadi and Pascale Bouchier, all members of the Vaccine Generation team, and Shessy Torres Morales from LUMC, dept of medical microbiology, for excellent scientific input and technical assistance. We would like to specially thank our colleagues of the Janssen Non-Clinical Safety (NCS) pathology laboratory in Beerse, Belgium, who have helped greatly with processing and analyzing all tissue samples for histological analyses.

## Author Contributions

J.vdL., S.R.H., E.vH., J.V., M.vH., M.K., E.S., L.dW., K.S., R.Z. and F.W. designed the experiments and analyzed the data, J.vdL., S.R.H., A.V., H.S., R.Z. and F.W. wrote the paper. L.D., L.vdF., L.R., J.L., D.B., R.Z. and F.W. contributed to the conception of the work. J.T., J.S. and L.M. contributed to the design of the experiments and performed the statistical analyses. E.vH., J.V., Y.C., M.B., K.F.dB., A.I.G., M.vH., T.D., S.M. and L.dW. performed the experiments and analyzed the data. All authors reviewed, critiqued, provided comments, and approved the text.

## Data Availability

All data that support the findings of this study are available from the corresponding author upon reasonable request.

