## Supplementary figures and tables for "Ad26.COV2.S-elicited immunity protects against G614 spike variant SARS-CoV-2 infection in Syrian hamsters and does not enhance respiratory disease in challenged animals with breakthrough infection after sub-optimal vaccine dosing"

### Fig S1

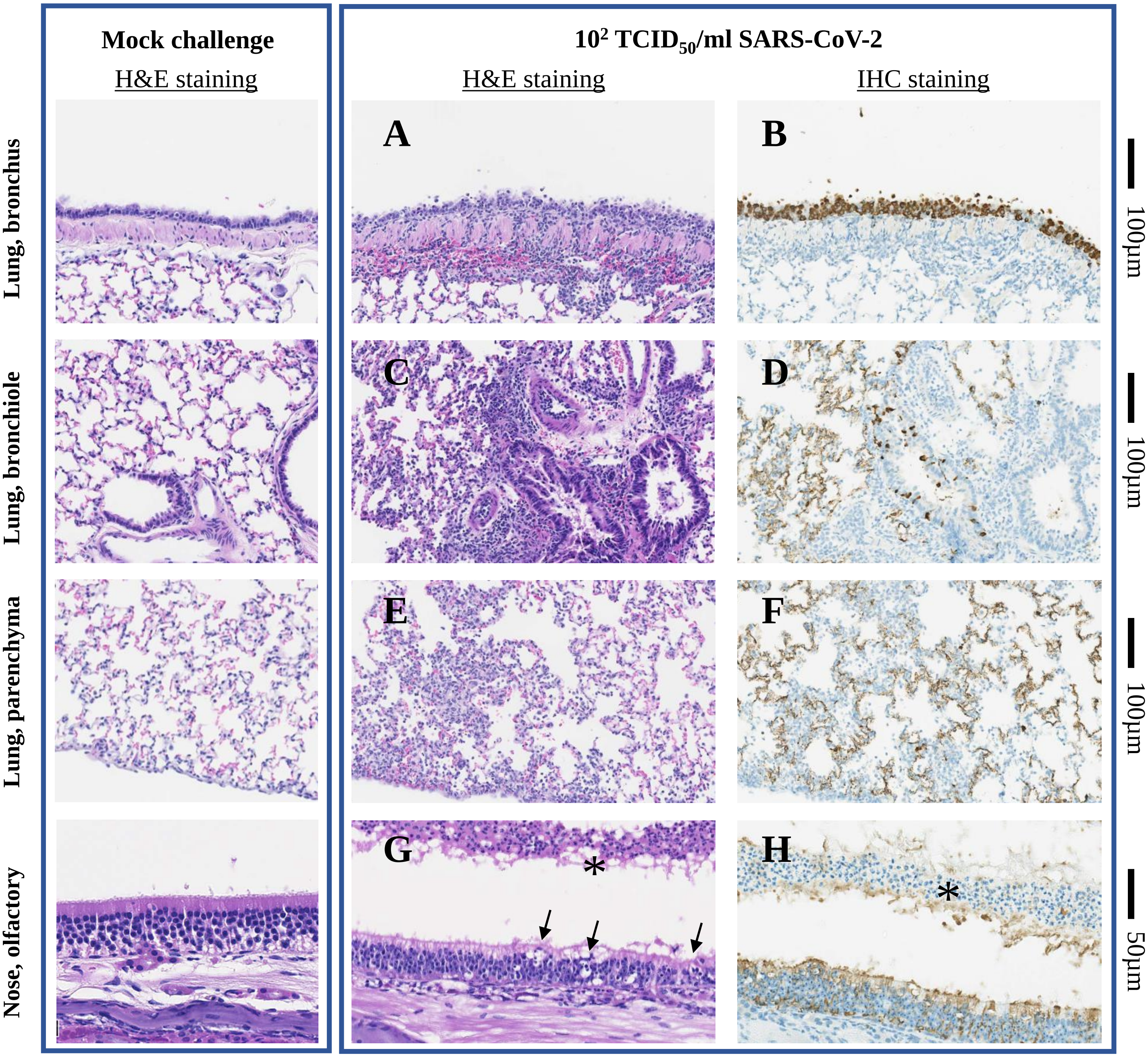


**Histopathologic features observed in hamster respiratory tissues.**

Syrian hamsters were inoculated intra-nasally with $10^2$ $TCID_{50}$ SARS-CoV-2 BetaCoV/Munich/BavPat1/2020, or mock-challenged with Vero E6 cell-supernatant. Four days post inoculation, hamsters were sacrificed, and left lung and nose tissues collected for histopathology.

**a)** Degeneration/necrosis of bronchial epithelium. Inflammation mononuclear and granulocytic lamina propria, with transmigration of granulocytes, Peribronchial cuffing, and peribronchial hemorrhages. **b)** SARS-CoV-2 NP positive bronchial epithelium cells. **c)** Degeneration/necrosis of bronchiolar epithelium, Peribronchiolar/perivascular cuffing and peribronchial hemorrhages. Infiltrate mononuclear/mixed cells in alveolar spaces/interstitium. **d)** SARS-CoV-2 NP positive bronchiolar epithelium cells, type I and II pneumocytes and some alveolar macrophages. **e)** Infiltrate mononuclear/mixed cells in alveolar spaces/interstitium, minimal hemorrhages. **f)** SARS-CoV-2 NP positive type I and II pneumocytes and some alveolar macrophages/mononuclear cells. **g)** Degeneration of olfactory epithelium (arrows) and presence of inflammatory cells (*) and sloughed cells in nasal cavity. **h)** SARS-CoV-2 NP positive olfactory epithelium and inflammatory cells/sloughed cells (*) in nasal cavity.

#### Fig S2:

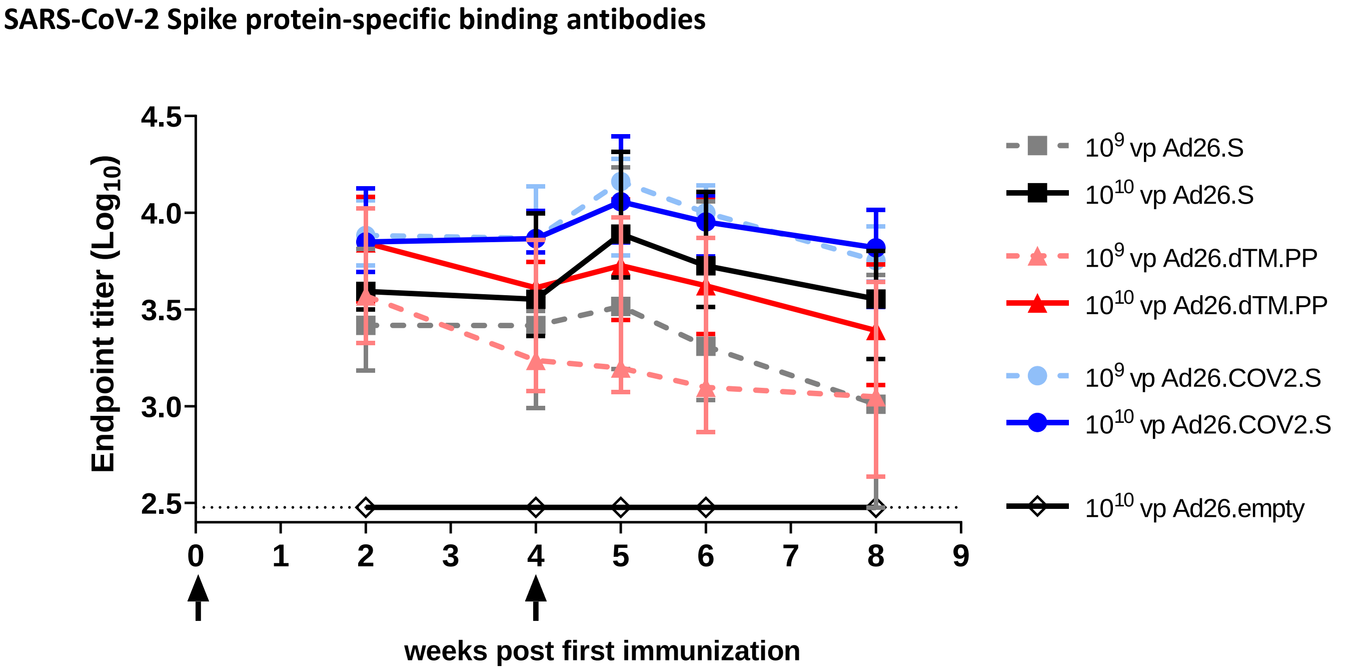


**SARS-CoV-2 spike protein binding antibody response, and ratio with neutralizing antibody responses, elicited by 1- and 2-dose Ad26 vaccine regimes in Syrian hamsters.** Syrian hamsters were immunized with either $10^9$ or $10^{10}$ VP of Ad26-based vaccine candidates Ad26.S, Ad26.dTM.PP or Ad26.COV2.S, or with $10^{10}$ vp of an Ad26 vector without gene insert as control (Ad26.empty). Four weeks after immunization half the hamsters per group received a second immunization with the same Ad26-based vaccine candidate (N=6 per group). Two and 4 weeks after one immunization, and 1, 2 and 4 weeks after the second immunization (weeks 5, 6 and 8 respectively), blood samples were collected and serum isolated for analysis of SARS-CoV-2 spike protein-specific antibody binding titers of hamsters receiving two immunizations (N=6 per group) by ELISA.

Dotted line indicates the lower limit of detection, median responses per group are indicated with horizontal lines. ELISA = Enzyme-linked Immunosorbent assay; wtVNA = wild-type virus neutralization assay; VP = virus particles.

**Fig S3**

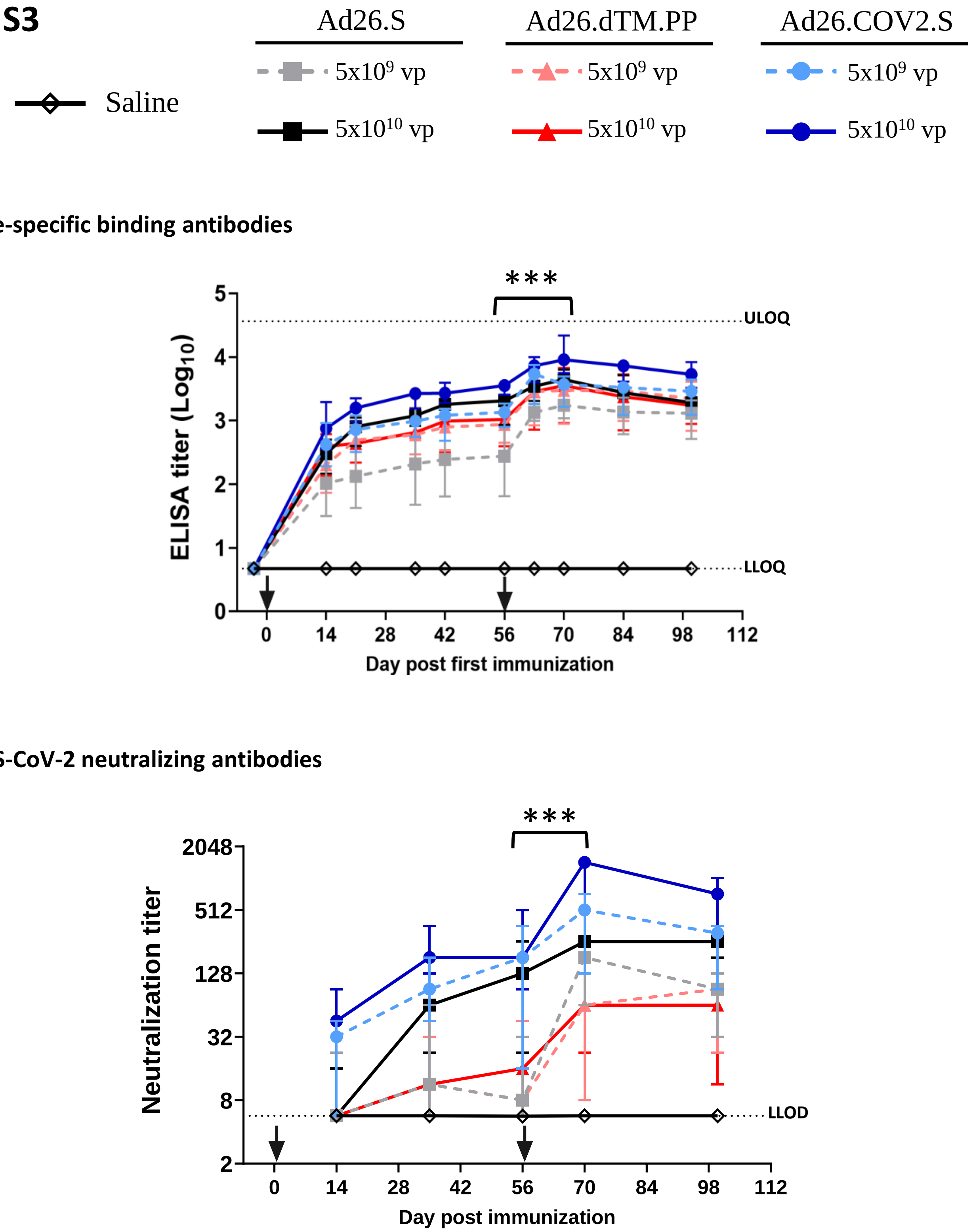


**SARS-CoV-2 spike protein binding antibody response and neutralizing antibody response elicited by a 2-dose Ad26 vaccine regime in New Zealand White rabbits.** New Zealand white rabbits were intramuscularly immunized with a 2-dose regimen of $5x10^9$ vp or $5x10^{10}$ vp Ad26.S, Ad26.dTM.PP, Ad26.COV2.S, or saline. Serum was sampled prior to immunization (day -3) and days 14, 21, 35, 42, 56, 63, 70, 84 and at sacrifice (days 99-101, depicted in the graph as day 100). **a)** SARS-CoV-2 spike protein-specific antibody binding titers were measured by ELISA. **b)** Neutralization titers were measured on days 14, 35, 56, 70 and at sacrifice.

Median responses per group are indicated with horizontal lines, vertical lines denote group ranges. Binding and neutralizing antibody titers of day 70 were compared with day 56 with a a-test from Tobit ANOVA, and asterisks indicated significantly higher titers induced by a second immunization with Ad26.S., Ad26.dTM.PP and Ad26.COV2.S: ***; $p<0.001$ . ELISA = Enzyme-linked Immunosorbent assay; wtVNA = wild-type virus neutralization assay; LLOQ = Lower Limit of Qualification; ULOQ = Upper Limit of Qualification; LLOD = Lower Limit of Detection; VP = virus particles; ANOVA = Analysis of Variance.

#### Fig S4

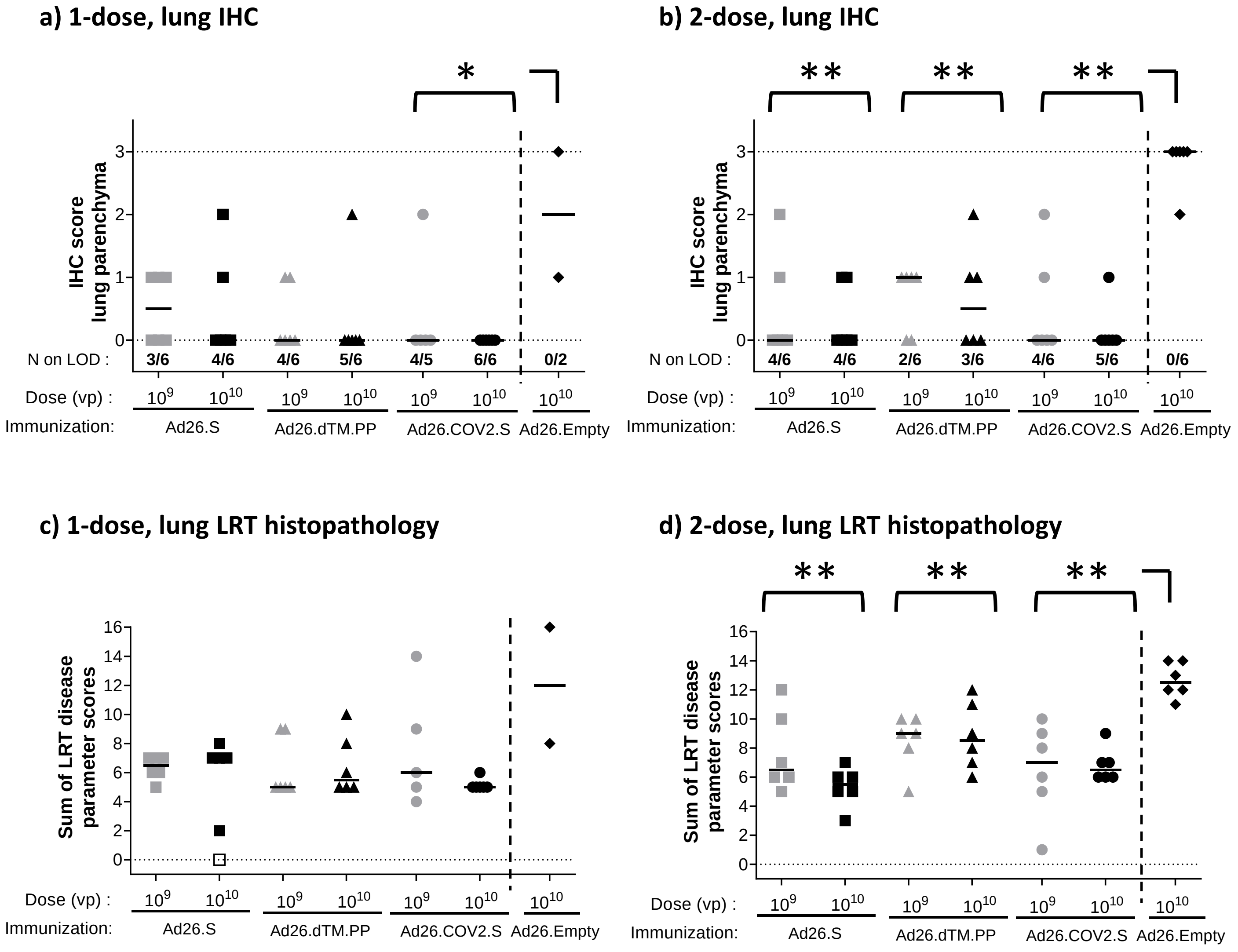


**Protection against SARS-CoV-2 IHC and histopathology in lung tissue of Syrian hamsters immunized with Ad26-based vaccine candidates.**

Syrian hamsters were intramuscularly immunized with a 1-dose regimen and a 2-dose regimen of Ad26.S, Ad26.dTM.PP, Ad26.COV2.S, or Ad26.empty (Ad26 vector not encoding any SARS-CoV-2 antigens). The hamsters received intranasal inoculation with $10^2$ $TCID_{50}$ SARS-CoV-2 strain BetaCoV/Munich/BavPat1/2020 4 weeks post-dose 1 (week 4), or 4 weeks post-dose 2 (week 8). Left lung tissue was isolated 4 days after inoculation for analysis of immunohistochemical SARS-CoV-NP staining and histopathology. Due to a technical failure, tissues of only 2 out of 8 hamsters immunized with one dose of Ad26.Emtpy could be analyzed. **a and b)** Presence of SARS-CoV-2 NP was determined by immunohistochemical staining. **c and d)** Lung tissue was scored for presence and severity of alveolitis, alveolar damage, alveolar edema, alveolar hemorrhage, type II pneumocyte hyperplasia, bronchitis, bronchiolitis, peribronchial and perivascular cuffing. Sum of scores are presented as sum of LRT disease parameters.

Median scores per group are indicated with horizontal lines. Dotted lines indicate the LLOD. Comparisons were performed between the vaccine groups across dose level, with the Ad26.Empty group by Mann-Whitney U-test, except for **panel c**. Statistical differences indicated by asterisks: *:p<0.05; **:p<0.01.

N on LOD = Number of hamsters on limit of detection, of the number of hamsters per group.; VP = virus particles; NP = Nucleocapsid protein; IHC = Immunohistochemistry; LRT = Lower Respiratory Tract.

#### Fig S5

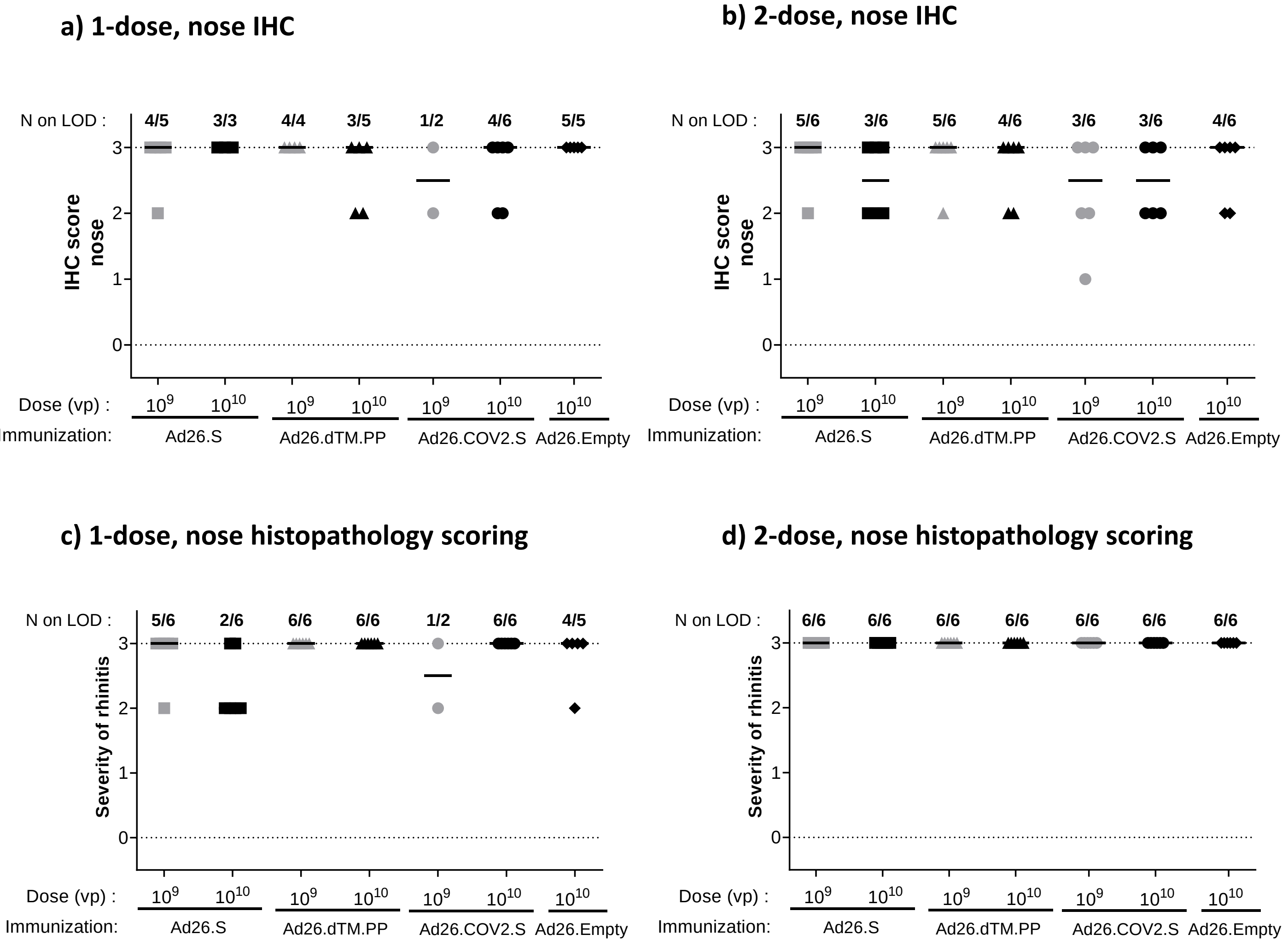


**Protection against SARS-CoV-2 IHC and histopathology in nasal tissue of Syrian hamsters immunized with Ad26-based vaccine candidates.**

Syrian hamsters were intramuscularly immunized with a 1-dose regimen and a 2-dose regimen of Ad26.S, Ad26.dTM.PP, Ad26.COV2.S, or Ad26.empty (Ad26 vector not encoding any SARS-CoV-2 antigens). The hamsters received intranasal inoculation with $10^2$ $TCID_{50}$ SARS-CoV-2 strain BetaCoV/Munich/BavPat1/2020 4 weeks post-dose 1 (week 4), or 4 weeks post-dose 2 (week 8). Left nasal tissue was isolated 4 days after inoculation for analysis of immunohistochemical SARS-CoV-NP staining and histopathology. Due to a technical failure, tissues of only 5 out of 8 hamsters immunized with one dose of Ad26.Emtpy could be analyzed. **a and b)** Presence of SARS-CoV-NP was determined by immunohistochemical staining. **c and d)** Nasal tissue was scored for severity of rhinitis.

Median scores per group are indicated with horizontal lines. Dotted lines indicate the LLOD. Comparisons were performed between the vaccine groups across dose level, with the Ad26.Empty group by Mann-Whitney U-test. Statistical differences indicated by asterisks: *:p<0.05; **:p<0.01.

LLOD = lower limit of detection; VP = virus particles; NP = Nucleocapsid protein; IHC = Immunohistochemistry. N on LOD = Number of hamsters on limit of detection, of the number of hamsters per group.

Table 1: Two-dose histopathology details

| Treatment : | Ad26.S | | Ad26.dTM.PP | | Ad26.COV2.S | | Ad26. Empty | N/A |
|---|---|---|---|---|---|---|---|---|
| Dose : | $10^9$ vp (N = 6) | $10^{10}$ vp (N = 6) | $10^9$ vp (N = 6) | $10^{10}$ vp (N = 6) | $10^9$ vp (N = 6) | $10^{10}$ vp (N = 6) | $10^{10}$ vp (N = 6) | N/A (N = 6) |
| Challenge : | SARS-CoV-2[a] | | | | | | | N/A[b] |
| Severity of alveolitis | **1.5** (1 - 2) | **1** (1 - 1) | **2** (1 - 2) | **1** (1 - 3) | **1** (0 - 3) | **1** (1 - 2) | **3** (2 - 3) | **0** (0 - 1) |
| Extent of alveolitis and/or alveolar damage | **1** (1 - 1) | **1** (1 - 1) | **1** (1 - 2) | **2** (1 - 2) | **1** (0 - 1) | **1** (1 - 2) | **1** (1 - 1) | **0** (0 - 1) |
| Extent plus severity of alveolitis | **2.5** (2 - 3) | **2** (2 - 2) | **3** (2 - 4) | **3** (3 - 4) | **2** (0 - 4) | **2** (2 - 3) | **4** (3 - 4) | **0** (0 - 2) |
| Thickening of alveolar septa (edema) | **0** (0 - 1) | **0** (0 - 0) | **0** (0 - 0) | **0** (0 - 1) | **0** (0 - 1) | **0** (0 - 0) | **1** (1 - 1) | **0** (0 - 0) |
| Alveolar hemorrhage | **0** (0 - 0) | **0** (0 - 0) | **0** (0 - 0) | **0** (0 - 0) | **0** (0 - 1) | **0** (0 - 0) | **1** (1 - 1) | **0** (0 - 0) |
| Type II pneumocyte hyperplasia | **1** (0 - 1) | **1** (0 - 1) | **1** (0 - 1) | **1** (0 - 1) | **1** (0 - 1) | **1** (0 - 1) | **1** (1 - 1) | **0** (0 - 1) |
| Severity of bronchitis | **1** (1 - 3) | **1** (1 - 2) | **3** (1 - 3) | **2.5** (1 - 3) | **1** (0 - 3) | **1** (1 - 3) | **3** (3 - 3) | **0** (0 - 0) |
| Severity of bronchiolitis | **1** (1 - 2) | **0.5** (0 - 1) | **1** (1 - 2) | **1** (1 - 2) | **1** (0 - 1) | **1** (1 - 1) | **1** (1 - 2) | **0** (0 - 0) |
| Degree of peribronchial and perivascular cuffing | **1** (1 - 2) | **1** (0 - 1) | **1** (1 - 1) | **1** (1 - 2) | **1** (1 - 1) | **1** (1 - 1) | **1.5** (1 - 2) | **0** (0 - 0) |
| Severity of tracheitis | **2** (1 - 3) | **1** (1 - 2) | **3** (2 - 3) | **3** (1 - 3) | **2** (0 - 3) | **1.5** (1 - 3) | **3** (2 - 3) | **0** (0 - 0) |
| Severity of rhinitis | **3** (3 - 3) | **3** (3 - 3) | **3** (3 - 3) | **3** (3 - 3) | **3** (3 - 3) | **3** (3 - 3) | **3** (3 - 3) | **0** (0 - 1) |

Median histopathology scores per group (range). Syrian hamsters were intramuscularly immunized with a 2-dose regimen with a 4-week interval, of $1{\times}10^9$ or $1{\times}10^{10}$ vp of Ad26.S, Ad26.dTM.PP, Ad26.COV2.S, or Ad26.Empty (Ad26 vector not encoding any SARS-CoV-2 antigens). N=6 per dose level per regimen. The hamsters received intranasal inoculation with $10^2$ $TCID_{50}$ SARS-CoV-2 strain BetaCoV/Munich/BavPat1/2020 4 weeks post Dose 2 (Day 56). A group of sentinel hamsters (N=6) received no immunization (Untreated) and were mock inoculated with Vero E6 cell supernatant. Lung, trachea and nasal turbinates were isolated at the end of the 4-day inoculation phase for histopathological analysis.

Histopathology scoring: Alveolitis, bronchitis, bronchiolitis, tracheitis and rhinitis severity: 0 = no inflammatory cells, 1 = minimal inflammatory cells, 2 = mild number of inflammatory cells, 3 = moderate inflammatory cells. Alveolitis extent, 0 = scattered, focal or focal/multifocal, 1 = multifocal <5%, 2 = multifocal 5-25%, 3 = >25%. Thickening of alveolar septa by (mononuclear/mixed cellular) infiltrate, alveolar haemorrhage and type II pneumocyte hyperplasia: 1 = minimal, 2= mild, 3 = moderate. Extent of peribronchial and perivascular cuffing, 0 = none, 1 = 1-2 cells thick, 2 = 3-10 cells thick, 3 = >10 cells thick.

[a] Inoculation with $10^2$ $TCID_{50}$ SARS-CoV-2 strain BetaCoV/Munich/BavPat1/2020 100 µL inoculum by the intranasal (IN) route.

[b] Mock-inoculation with Vero E6 cell supernatant, 100 µL by the intranasal (IN) route.

Table. N = number of animals per group; vp = virus particles.

Table 2: Ad26.COV2.S dose titration histopathology details

| Treatment : | Ad26.COV2.S | | | | Ad26.Irr | |
|---|---|---|---|---|---|---|
| Dose : | $10^7$ vp (N = 8) | $10^8$ vp (N = 8) | $10^9$ vp (N = 8) | $10^{10}$ vp (N = 8) | $10^{10}$ vp (N = 8) | $10^{10}$ vp (N = 8) |
| Challenge : | SARS-CoV-2 | | | | | Mock |
| Severity of alveolitis | **1** (1 - 2) | **1** (1 - 2) | **0.5** (0 - 1) | **1** (0 - 1) | **1.5** (1 - 3) | **0** (0 - 1) |
| Extent of alveolitis and/or alveolar damage | **1** (0 - 2) | **0** (0 - 1) | **0** (0 - 0) | **0** (0 - 0) | **0.5** (0 - 2) | **0** (0 - 0) |
| Extent plus severity of alveolitis | 2 (1 - 4) | **1** (1 - 3) | **0.5** (0 - 1) | **1** (0 - 1) | **2** (1 - 5) | **0** (0 - 1) |
| Thickening of alveolar septa (edema) | 0.5 (0 - 3) | **0** (0 - 1) | **0** (0 - 0) | **0** (0 - 0) | **0.5** (0 - 2) | **0** (0 - 0) |
| Alveolar haemorrhage | **0** (0 - 2) | **0** (0 - 3) | **0** (0 - 2) | **0** (0 - 2) | **2** (0 - 3) | **0** (0 - 2) |
| Type II pneumocyte hyperplasia | **0** (0 - 0) | **0** (0 - 0) | **0** (0 - 0) | **0** (0 - 0) | **0** (0 - 1) | **0** (0 - 0) |
| Severity of bronchitis | **2** (0 - 3) | **1** (0 - 3) | **0** (0 - 0) | **0** (0 - 0) | **3** (2 - 3) | **0** (0 - 0) |
| Severity of bronchiolitis | **0** (0 - 3) | **0** (0 - 3) | **0** (0 - 0) | **0** (0 - 1) | **2** (0 - 3) | **0** (0 - 0) |
| Degree of peribronchial and perivascular cuffing | **0.5** (0 - 2) | **1** (0 - 2) | **0** (0 - 0) | **0** (0 - 1) | **2** (1 - 2) | **0** (0 - 1) |
| Severity of rhinitis | **3** (2 - 3) | **2** (0 - 3) | **3** (0 - 3) | **2** (1 - 3) | **3** (1 - 3) | **0** (0 - 0) |
| Severity of tracheitis | **2** (1 - 3) | **1** (0 - 3) | **0.5** (0 - 2) | **0** (0 - 3) | **2** (1 - 3) | **0** (0 - 0) |

Median histopathology scores per group (range). Syrian hamsters were intramuscularly immunized with $10^7$, $10^8$, $10^9$ or $10^{10}$ vp of Ad26.COV2.S, N=8 per group, or $10^{10}$ vp Ad26.Irr (Ad26 vector not encoding any SARS-CoV-2 antigens, N=16). The hamsters received intranasal inoculation with $10^2$ $TCID_{50}$ SARS-CoV-2 strain BetaCoV/Munich/BavPat1/2020 4 weeks post immunization (Day 28). Eight hamsters which were immunized with Ad26.RSV.gLuc were mock inoculated with Vero E6 cell supernatant (Mock). Lung, trachea and nasal turbinates were isolated at the end of the 4-day inoculation phase for histopathological analysis.
Histopathology score: Alveolitis, bronchitis, bronchiolitis, tracheitis and rhinitis severity: 0 = no inflammatory cells, 1 = minimal inflammatory cells, 2 = mild number of inflammatory cells, 3 = moderate inflammatory cells. Alveolitis extent, 0 = scattered, focal or focal/multifocal, 1 = multifocal <5%, 2 = multifocal 5-25%, 3 = >25%. Thickening of alveolar septa by (mononuclear/mixed cellular) infiltrate, alveolar haemorrhage and type II pneumocyte hyperplasia: 1 = minimal, 2= mild, 3 = moderate. Extent of peribronchial and perivascular cuffing, 0 = none, 1 = 1-2 cells thick, 2 = 3-10 cells thick, 3 = >10 cells thick.
Ad26.Irr = Ad26 vector not encoding any SARS-CoV-2 antigens; N = number of animals per group; vp = virus particles.
